# The long non-coding RNA *FAM30A* regulates the Musashi2-RUNX1 axis and is required for LSC function in AML cells

**DOI:** 10.1101/2024.10.13.618058

**Authors:** Jaime Calvo Sánchez, Mary T. Scott, Athina Varnava, Tala Alakhras, Alice Wedler, Danny Misiak, Nadine Bley, Christian Ihling, Andrea Sinz, Karen Keeshan, Stefan Hüttelmaier, David Vetrie, Marcel Köhn

## Abstract

High expression of the long non-coding RNA (lncRNA) *FAM30A* has been previously associated with leukemic stem cell (LSC) activity and poor prognosis in both adult and paediatric acute myeloid leukaemia (AML) patients, yet it has not been functionally studied. This study provides the first cellular characterization of *FAM30A* focussing on an internal tandemly organised region, referred to as *FAM30A* repeats. *FAM30A* levels correlated with canonical AML LSC signatures and *FAM30A* depletion decreased cell viability as well as increased sensitivity to chemotherapeutics. It also inhibited colony formation, promoted granulocytic differentiation and abrogated leukemic engraftment in murine bone marrow *in vivo*. Overexpression of *FAM30A* repeats in this setting enhanced stemness, proliferation, chemoresistance, and engraftment thus highlighting the biological relevance of this region for LSC biology. On the molecular level, *FAM30A* repeats interact with the pro-LSC regulator Musashi-2 (MSI2), positively influencing expression of its targets including RUNX1 isoforms. We herein uncover that this *FAM30A*-MSI2-RUNX1 regulatory loop is of potential relevance for LSC maintenance in AML. These findings provide valuable insights into *FAM30A*’s cellular role and highlight its targeting potential for eliminating LSCs and improving treatment outcomes in AML patients.

## INTRODUCTION

A significant challenge of acute myeloid leukaemia (AML) patients is the poor prognosis and high relapse rate following aggressive chemotherapy regimens. Disease relapse is believed to be driven mainly by leukaemia stem cells (LSCs) which are typically resistant to standard-of-care treatments and persist within AML patients, undergo clonal diversification and ultimately lead to more aggressive forms of the disease^1, 2^. Deciphering the expression and functionality of LSC-associated genes has marked a pivotal breakthrough in AML research as it enables the identification and targeting of LSCs, potentially driving the development of novel therapeutics for AML patients^3, 4^.

Long non-coding RNAs (lncRNAs) constitute a class of transcripts (>200 nucleotides) which are increasingly investigated, especially with the emergence of advanced sequencing techniques^5^. Growing evidence highlights how individual lncRNAs can affect cell fate decisions within the haematopoietic hierarchy, where their dysregulation may contribute to the onset of haematological malignancies, including AML^6–8^. In fact, the role of lncRNAs as important prognostic markers and potential therapeutic targets in AML is rapidly expanding and aberrant expression of certain lncRNAs has been associated with worse clinical outcome^9^ and AML-specific recurrent mutations^10^. Moreover, high expression of certain lncRNAs was significantly correlated with well-established LSC signatures^11^ even though only some of them were proven to alter LSC activity *in vitro* and *in vivo*^7, 11, 12^. For example, the cytoplasmic stem-cell specific lncRNA *HOXA10-AS* was demonstrated to act *in trans* to induce the transcription of NF-kB target genes and promote leukemogenesis by blocking monocytic differentiation in KMT2A-rearranged AML^13^.

Among these prognostic high-risk genes, the putative lncRNA *FAM30A* (also named as *KIAA0125*) is the most consistently upregulated lncRNA in an LSC^high^-associated lncRNA signature in young and older AML patients^11^. Apart from being included in other lncRNA signatures for AML^14, 15^, *FAM30A* has consistently been included in several other independent gene signatures linked to AML LSCs activity and poor prognosis in both adult^16–18^ and paediatric AML^19, 20^. The biochemical and molecular characteristics of *FAM30A* transcripts have not been investigated so far and RNA features that mediate its cellular roles especially in cancer are unknown. While *FAM30A* has been proposed to have context-specific tumour suppressive^21–23^ or oncogenic potential^24–26^ in different cell types^27^, the underlying molecular role whereby *FAM30A* exerts its function in AML LSCs still remains unexplored. Considering the high expression of *FAM30A* in functionally validated AML LSCs fractions when compared to leukemic blasts^16^, along with the marked upregulation of *FAM30A* in LSCs after induction chemotherapy^28^, we hypothesized that altering *FAM30A* expression could have an impact on LSC features and leukemogenesis. In this study we aimed to dissect the significance of *FAM30A* in LSC biology in AML cells. We identified the protein interactors of a tandemly repeated region within the *FAM30A* sequence and described its functional role in regulating LSC features.

## RESULTS

### *FAM30A* expression in AML and its association with high LSC content

*FAM30A* levels were first analysed across bone marrow (BM) samples of acute and chronic leukaemia subclasses extracted from the Microarray Innovations in LEukemia (MILE) multi-center study^29^. The results showed that *FAM30A* levels were significantly higher in AML samples (n=516, P=4.93E-14) and in other leukaemia types compared to healthy BM (n=71, Figure 1A) as well as solid tumour types (TCGA) (Supplementary Figure 1A). Aiming to identify putative lncRNAs with oncogenic potential in AML samples from The Cancer Genome Atlas (TCGA) vs healthy BM (GTEx) in an unbiased manner, differential gene expression data was extracted from GEPIA 2^30^. The analysis (cut-off: log_2_Fold-change > 1, P < 0.01) showed that *FAM30A* was the third most significantly upregulated lncRNA (51-fold change) when compared to healthy BM samples (Figure 1B). In line with previous studies and the proposed oncogenic potential of *FAM30A* in AML, high *FAM30A* levels are strongly correlated with shortened overall survival in AML patients from different cohorts (n=1608) when compared to patients with low *FAM30A* expression (9.1 vs 20.1 months, Figure 1C)^14,31–34^. *FAM30A* was highly expressed in patients with more aggressive karyotypes and specific mutational status associated with poor prognosis (i.e. *FLT3-ITD*, *EVI1* mutations)^35^ (Supplementary Figure 1B-D). On the other hand, expression of *FAM30A* was lower in AML patients with better prognosis (i.e. t(8;21) and t(9;11)), in line with results from a previous study^25^. Moreover, significantly higher *FAM30A* levels were seen in AML subtypes with more stem-cell like and higher chemoresistance properties (M0 and M1)^36^ compared to others subtypes based on the French-American-British (FAB) system (Figure 1D). Next, we screened publicly available transcriptomic data for *FAM30A* expression from functionally validated LSC fractions (LSC+) with varying CD34/CD38 status and compared them to non-LSC enriched fractions (LSC-, total n=78)^16^. This revealed that *FAM30A* expression was significantly higher in the LSC-enriched fraction and the highest in the CD34+CD38- compartment (4.5 times more *FAM30A* than in the CD34-fraction). This fraction was also shown to be where the LSC frequency *in vivo*, long-term repopulating and chemoresistance capacities are the highest (Figures 1E, F)^37^. Utilizing recent single-cell RNA-sequencing (scRNA-seq) data derived from BM samples from healthy donors (n=10), adult (n=20) and paediatric (n=22) AML patients^38^, the analysis showed that *FAM30A* expression was restricted to HSC-like blasts (HSC) and immature progenitors, myeloid-committed (i.e. MPP, MEP) with lower apparent expression in their differentiated progenies (Figure 1G). This pattern of *FAM30A* expression followed a pattern similar to other well-known LSC-associated and highly *FAM30A*-correlated genes such as CD34 and HOPX (Supplementary Figure 1E)^28^.

**Figure 1.**
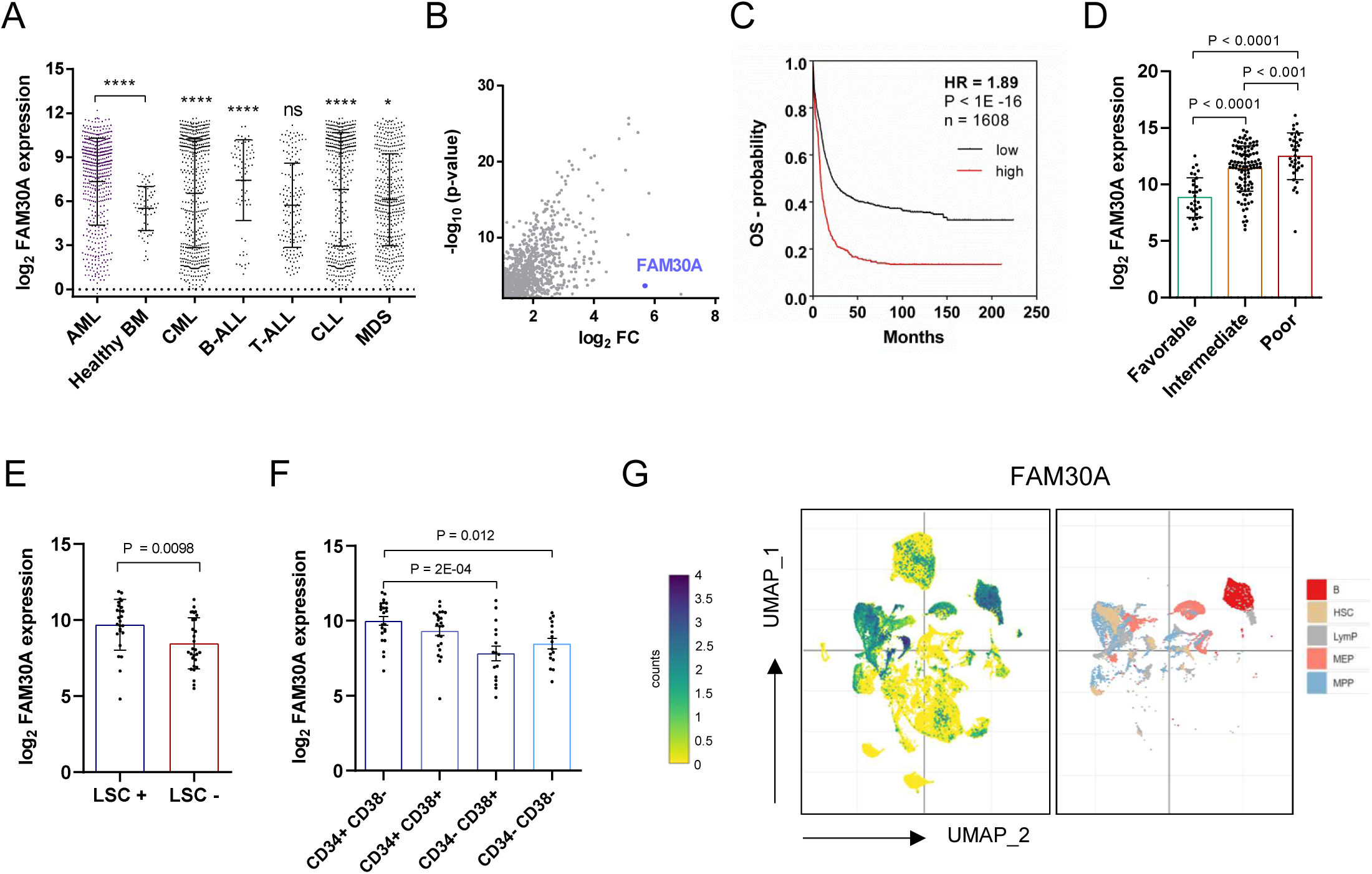
*FAM30A* levels correlate with high LSC content and worse prognosis in AML patients. **A.** *FAM30A* levels (log_2_ RNA expression) extracted from the MILE study in patients with different leukaemia subtypes (AML, n=516; healthy BM, n=70; CML, n=497; B-ALL, n=80; T-ALL, n=165; CLL, n=428; MDS, n=426). Statistical significance was calculated using healthy BM as control. **B.** Representative graph of upregulated lncRNAs (cut-off: log_2_Fold Change (FC)>1, P < 0.01) in AML samples (TCGA-LAML, PanCancerAtlas, n=173) compared to healthy BM samples (GTEX). **C.** Kaplan-Meier survival plot of the overall survival of AML patients (n=1608), categorized according to *FAM30A* gene expression (high vs low, based on median expression). **D.** *FAM30A* expression data (log_2_ RNA expression) extracted from TCGA (PanCancerAtlas) with each karyotype assigned to be favourable (n=32), intermediate (n=105) or poor (n=36) prognosis according to current guidelines. **E.** *FAM30A* levels extracted from validated LSC (n=29) and non-LSC (n=25) fractions in AML patients^16^. **F.** *FAM30A* expression data derived from validated fractions from E. with varying CD34, CD38 surface expression. **G.** UMAP visualization map depicting normalised counts of *FAM30A* (left panel) clusters based on cell-type identity (right panel) using the scRNAseq portal (Broad Institute). Data extracted from^38^. Statistical significance was calculated using unpaired two-tailed student t-test (* P < 0.05, **** P < 0.0001, ns – not significant). Data is displayed as mean values and error bars represent SEM. AML: acute myeloid leukaemia, BM: bone marrow, CML: chronic myeloid leukaemia, B-ALL: B-cell acute lymphoblastic leukaemia, T-ALL: T-cell acute lymphoblastic leukaemia, CLL: chronic lymphoblastic leukaemia, MDS: myelodysplastic syndrome, HR: hazard ratio.

### Cellular characterization of *FAM30A*

After a comprehensive analysis of the *FAM30A* sequence (~10kb), we identified a repetitive region at the end of exon 5, which we hereafter named the *FAM30A* repeat region. This region contains six tandemly repeated sequences, two of 76 bp and four of 78 bp, amounting to 464 bp in total (Figure 2A). After multiple sequence alignment of the repeats we observed that the nucleotide identity among the six repeats was very high and prompted us to investigate whether this region along with other flanking parts of the RNA sequence is evolutionary conserved. *FAM30A* transcripts can only be found in primates and are not detectable on genomic level in other mammals like rodents. Furthermore, an in-depth genomic analysis of primate genomes suggested that humans display the highest number of repeats, with the start of this repeat region showing the highest conservation score upon alignment of primate genomes (Supplementary Figure 2A, B). This underscores the relevance of investigating the molecular function of *FAM30A* repeats as there is a potential evolutionary pressure to increase the number of *FAM30A* repeats. Moreover, *FAM30A* exhibited the highest expression levels in HSCs and other myeloid precursors (i.e. CMP, GMP) as well as its expression was confined to immune-related tissues, following a pattern consistent with our earlier findings (Figure 2B, Supplementary Figure 2C). RT-qPCR results across a panel of AML cell lines showed that *FAM30A* was highly expressed in the CD34+ AML cell lines KG-1a and GDM-1 (Figure 2C). In order to determine the exon usage of *FAM30A* in KG-1a cells we analysed RNA-sequencing data and observed that mainly exon 1 to 5 are expressed (Exon 2-6 based on NR_026800.2, Supplementary Figure 2D). Detailed analyses of *FAM30A* exon usage via RT-qPCR confirmed that exon 5 was constitutively expressed in KG-1a cells (Supplementary Figure 2D).

**Figure 2.**
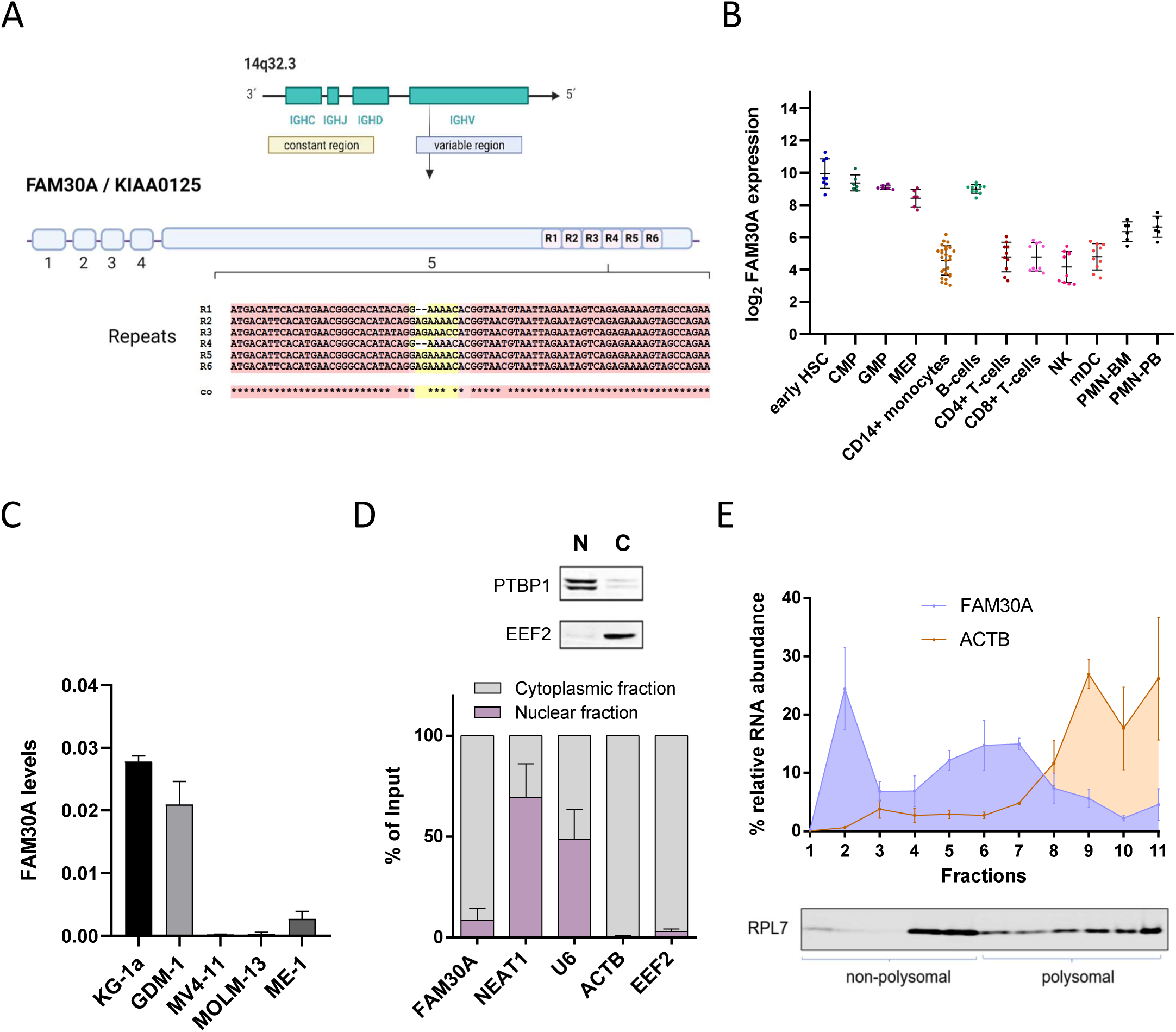
Cellular characterization of the cytoplasmic lncRNA *FAM30A*. **A.** Schematic of the genomic locus around *FAM30A* at the end of human chromosome 14, which is antisense to the *IGHV* locus. Lower panel depicts a detailed illustration of *FAM30A* sequence and the high conservation score (in red) among the six tandem repeats (R1-6). **B.** *FAM30A* expression levels extracted from^74^ in different haematopoietic cell types. **C.** Relative *FAM30A* RNA levels after RT-qPCR analysis in AML cell lines, normalized to *ACTB*. **D.** Confirmation of subcellular fractionation via Western blot by detecting PTBP1 in the nuclear fraction and EEF2 in the cytoplasm (higher panel). Nuclear versus cytoplasmic localization of *FAM30A*, *NEAT1, U6* RNA and *ACTB, EEF2* mRNA in KG-1a cells (lower panel). **E.** Targeted profiling of relative *ACTB* and *FAM30A* RNA abundances in KG-1a cells in non-polysome associated (1-6) and polysomal fractions (7-11) isolated after sucrose based gradient centrifugations. Representative western blot of RPL7 is shown as loading control for identification of fractions. Data is displayed as mean values and error bars represent SEM.

Since the subcellular localization may be connected to the biological function of an lncRNA^39^, digitonin-based subcellular fractionation was performed in KG-1a cells with fraction-specific controls *NEAT1, U6* (nuclear ncRNAs) and *ACTB*, *EEF2* (cytoplasmic mRNAs). As a result, we observed that *FAM30A* was predominantly located in the cytoplasm (more than 90% of the total transcript, Figure 2D). To determine whether *FAM30A* has protein-coding potential, we used sucrose gradient-based fractionation followed by targeted polysome profiling in KG-1a cells. We determined the abundances of *ACTB* and *FAM30A* RNAs on the isolated fractions and these were compared to ribosomal protein 7 (RPL7) distribution. As expected, *ACTB* mRNA as an example for a highly translated mRNA was enriched in the low and high molecular weight polysomal fractions whereas *FAM30A* RNA was enriched in only small complexes as well as slightly larger complexes co-migrating with monosomal fractions (70% of the total transcript). This indicates that *FAM30A* is a cytoplasmic lncRNA that does not associate strongly with polysomes and is therefore not efficiently translated at steady-state conditions in KG-1a cells (Figure 2E).

### Deregulation of *FAM30A* and its repeats alter proliferation, chemoresistance and myeloid differentiation in AML cells

To explore the oncogenic role of *FAM30A* and its repeats in AML, we engineered fluorescently tagged control cells (CTRL), *FAM30A* knockdown cells (KD), *FAM30A* repeat overexpression cells (OE) and a recovery cell line combining the KD with the OE (REC) in the KG-1a background (Figure 3A). We overexpressed the repeats only, since our conservation studies already suggested the biological relevance of this transcript region. Total RNA was isolated from these cell lines and subjected to bulk RNA sequencing which confirmed silencing of endogenous *FAM30*A and the overexpression of the repeats within the respective samples (Figure 3B). Gene Ontology (GO) and gene set enrichment analyses (GSEA) showed that gene expression patterns from the KD cells significantly correlated with downregulation of canonical AML LSC signatures (CE_HSC_LSC, LSC_R)^16^ and pro-survival pathways important for LSC maintenance (FDR < 0.001, MSigDB, Hallmarks) (Figure 3C, D; Supplementary Table S7 and S8).

**Figure 3.**
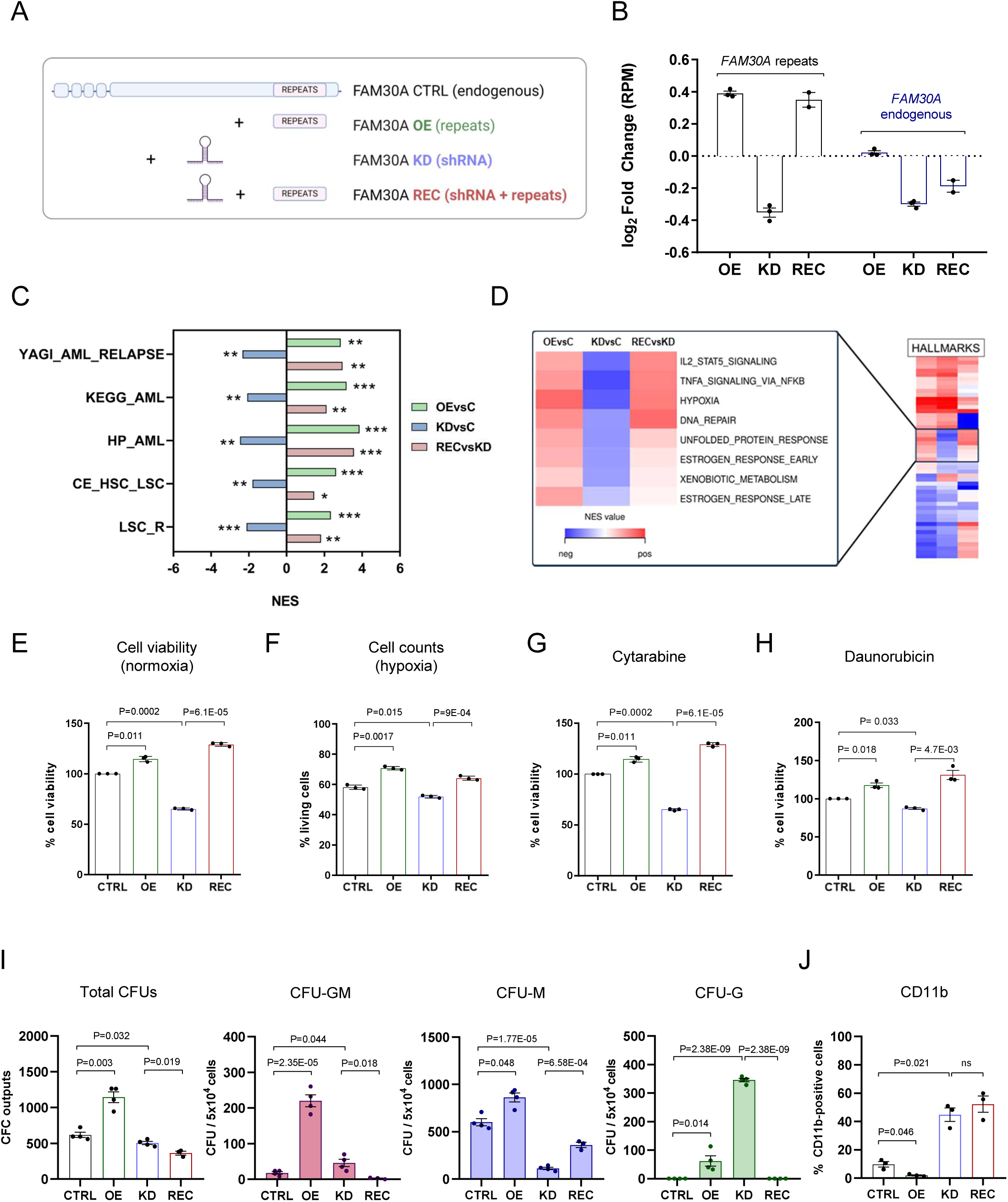
Modulation of *FAM30A* expression affects LSC features in AML cells. **A.** Illustration of the experimental design used to perform knockdown and overexpression studies of *FAM30A* in AML cells. **B.** Quantitative analysis derived from RNA-seq performed in stable KG-1a cell lines. Data is expressed in terms of log_2_ FC (RPM) of *FAM30A* repeat and its flanking region (*FAM30A* endogenous), normalised to control cells. **C.** GSEA analysis (C2, MSigDB) derived from RNA-seq in stable KG-1a cell lines. **D.** Heatmap of GSEA analysis (Hallmarks, MSigDB) derived from RNA-seq as in C. **E-G**. Comparison of viable cells/cell counts of stable KG-1a cell lines after 48 hours incubation in normoxic (E) or hypoxic (F) conditions and treatment with cytarabine (3 µM) (G) and daunorubicin (0.1 µM) (H). **I.** From left to right: quantification of number of colonies of KG-1a cell lines cultured in methylcellulose medium for 10 days, expressed in total colony-forming cell outputs (CFC) or colony-forming units for granulocytes/macrophages (CFU-GM, CFU-M and CFU-G). **J.** Quantification after FACS analysis of CD11b-positive cells in stable KG-1a cell lines in liquid culture for 5 days. Statistical significance was calculated using unpaired two-tailed student t-test (* P < 0.05, ** P < 0.01, *** P < 0.001). Data is displayed as mean values and error bars represent SEM. RPM: reads per million; NES: normalised enrichment score. CFU-GM: colony forming unit-granulocyte-macrophage; CFU-M: colony forming unit macrophage; CFU-G: colony forming unit-granulocyte.

We cultured the four KG-1a cell lines under normal growth conditions for two days and observed that repeat overexpression significantly increased cell viability compared to CTRL whereas the KD cells exhibited decreased cell viability which was recovered upon REC conditions (Figure 3E). As hypoxia was one of the pathways correlated to *FAM30A* levels (Figure 3D), we determined how the cells react to these conditions. The cell counts revealed that OE/REC cells again had a growth advantage even under hypoxic conditions (Figure 3F). We also determined how KG-1a stable cell lines behave under severe apoptotic conditions. OE/REC cells showed increased cell viability as well as decreased apoptosis with inverse behaviour of KD cells after treatment with cytarabine and daunorubicin (IC_50_ = 3 µM and 0.5 µM, respectively) compared to control (Figure 3G, H; Supplementary Figure 3A-C). The increased chemoresistance upon overexpression of *FAM30A* repeats was also confirmed in the *FAM30A*^low^ AML cell lines MV4-11 and ME-1 (Supplementary Figure 3D).

Decreased total CFC outputs (>CFU-G) and increased lineage commitment (>CD11b expression) were also observed in the *FAM30A* KD cells (Figure 3I, J; Supplementary Figure 3E). Most of these phenotypic changes were rescued in the REC cell line. Overexpression of *FAM30A* repeats (OE) resulted in increased total CFC outputs (>CFU-GM) compared to control, while lineage commitment was reduced (<CD11b).

### *FAM30A* repeats interact and modulate Musashi-2 protein levels (MSI-2) in AML cells

In order to investigate the molecular function of the *FAM30A* repeats, we aimed at identifying its interacting partners in AML cells. Thus, we generated biotin-labelled *FAM30A* repeat RNAs for RNA pull-down assays which were incubated with KG-1a cell lysates. This was followed by quantitative proteomics (LC-MS/MS analysis) for an unbiased approach to identify associated RNA-binding proteins (RBPs). We identified several RBPs with high reliability to interact with the *FAM30A* repeat, including some reportedly associated with AML-leukemogenesis e.g. MSI2, HNRNPA1, TNIK, SYNCRIP or DDX21 (Figure 4A, complete list provided in Supplementary Table S9). Functional clustering analysis (GO) of the significantly enriched proteins revealed an association with biological processes connected to mRNA translation (Figure 4B). *In silico* screening (regRNA2.0, RBPmap) to identify putative RBP motifs across *FAM30A* sequence revealed three conserved MSI2-binding sites, characterized by UAG-containing motifs^40^ located within each single *FAM30A* repeat (Figure 4C). We focused on MSI2 (p-value 9.57E-06, log_2_FC=3.57) since it is a well-known cytoplasmic pro-LSC regulator mediating mRNA translational control in AML^41, 42^ whose expression also highly correlates with *FAM30A* in AML patients (TCGA PanCancerAtlas cohort, Supplementary Figure 4A). Additionally, scRNAseq data from AML patients showed that the expression pattern of *FAM30A* and MSI2 is highly correlated in HSCs and immature myeloid progenitors (Supplementary Figure 4B). The association between *FAM30A* repeats and MSI2 was validated via Western blot analysis, RNA-immunoprecipitation (RIP) and available MSI2-CLIP data in *FAM30A*^high^ AML cells. We also tested the specificity of the *FAM30A*-MSI2 interaction by mutating the three MSI2 motifs which abrogated association with MSI2 (Figure 4C, D; Supplementary Figure 4C). Moreover, high expression of *FAM30A* repeats was correlated with higher MSI2 protein abundances that influenced protein levels of direct MSI2-mRNA targets (i.e. SMAD3)^42^ (Figure 4E, F). These findings revealed the association of the lncRNA *FAM30A* with MSI2, highlighting their potential as modulatory circuit in AML pathogenesis.

**Figure 4.**
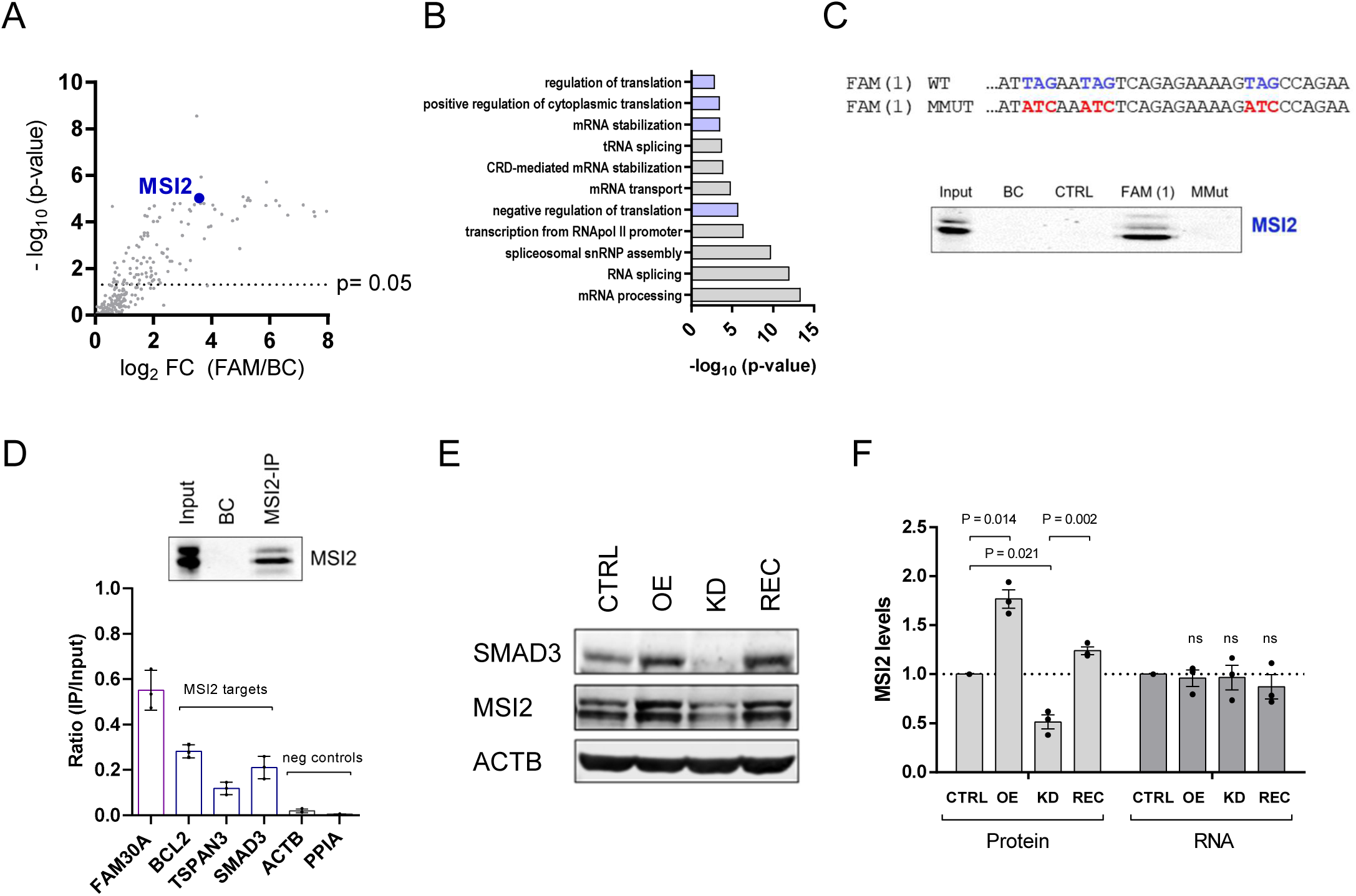
MSI2 interacts with *FAM30A* repeats in AML cells. **A.** Plot depicting enriched proteins after RNA-pulldown followed by LC-MS/MS analysis using biotinylated *FAM30A* repeats (FAM) as bait vs bead control (BC) in KG-1a cells. **B.** Functional clustering analysis (GO) for significantly enriched proteins FAM/BC (P< 0.05). Highlighted in blue are biological processes associated with MSI2 protein. **C.** Schematic of the probes used for RNA pulldown: a single wild-type *FAM30A* repeat (FAM(1) WT) with the three MSI2-binding sites (highlighted in blue), and a *FAM30A* repeat (FAM (1) MMUT) with mutated MSI2-binding sites (in red). Beads only (BC) as well as a control RNA neighbouring the *FAM30A* repeats (CTRL) were used as specificity controls. A representative Western blot analysis for MSI2 association in RNA pulldowns is shown. **D.** Western blot analysis for validation of immunoprecipitated endogenous MSI2 and RT-qPCR analysis of MSI2-associated RNAs (lower) in KG-1a cells. **E.** Representative Western blot of stable KG-1a cell lines for MSI2 protein and its validated target SMAD3. ACTB was used as a loading control **F.** Quantitative analysis of Western blot (light grey) RT-qPCR analysis (dark grey) for MSI2, harvested in cells from E. Statistical significance was calculated using unpaired two-tailed student t-test. Data is displayed as mean values and error bars represent SEM.

### The role of *FAM30A* repeats on RUNX1 transcriptional activation via MSI2

To date, the connection of *FAM30A* and RUNX1 has not been experimentally investigated in AML. A previous study predicted *in silico* the possibility of RUNX1 directly regulating the transcription of the *FAM30A* gene^25^. GSEA analyses (MsigDB) performed in KG-1a stable cell lines showed a highly significant correlation between expression of *FAM30A* repeats and transcriptional regulation by RUNX1 (a known regulator of LSCs maintenance)^43^ and its downstream targets (Figure 5A). Since these changes in RUNX1 activity were driven by the MSI2-interacting region of *FAM30A,* we sought to investigate whether MSI2 might regulate RUNX1. Analysis of publicly available MSI2 CLIP-seq data from AML and CML cells^42, 44^ revealed that MSI2 differentially binds to the 5’ and 3’ untranslated regions (UTR) of the three primary RUNX1 isoforms (Supplementary Figure 5A). To further validate this finding, we performed immunoprecipitation of MSI2 followed by detection of bound transcripts by RT-qPCR (RIP). This set of RIP experiments confirmed that MSI2 is strongly associated with the RUNX1C isoform, while the RUNX1A/B isoforms were not as strongly enriched (Figure 5B). To test the consequence of this binding behaviour we analysed RUNX1 protein levels upon *FAM30A* modulation. Here, the *FAM30A* repeat overexpression led to a significant increase of all protein isoforms of RUNX1. However, in the *FAM30A* KD setting the RUNX1 isoforms were differentially regulated (RUNX1C decreased, RUNX1A/B increased), an effect which was largely recovered under REC conditions (Figure 5C; Supplementary Figure 5B). Interestingly, the RUNX1 mRNA levels remained essentially unchanged upon *FAM30A* modulation (Supplementary Figure 5C), suggesting that regulation of RUNX1 by *FAM30A* occurred primarily at the protein level. Moreover, these changes in RUNX1C/RUNX1 protein levels were also correlated with MSI2 protein levels (Figure 5C). Consistently, we also observed the concomitant upregulation of RUNX1 isoforms as well as MSI2 in the FAM30^low^ AML cell line MV4-11 (Supplementary Figure 5D). To analyse RUNX1 transcriptional activity we utilized K562 cells stably overexpressing *FAM30A* repeats in an MSI2 knockdown setting. These cells were transfected with luciferase reporter constructs comprising RUNX1-binding sites (two or three) to assess transcriptional activity. These reporter analyses confirmed decreased RUNX1 activity upon MSI2 depletion as well as increased RUNX1 activity upon *FAM30A* repeat overexpression (Figure 5D, E). Hence, we confirmed *in cellulo* that RUNX1 activity is influenced by both *FAM30A* and MSI2.

**Figure 5.**
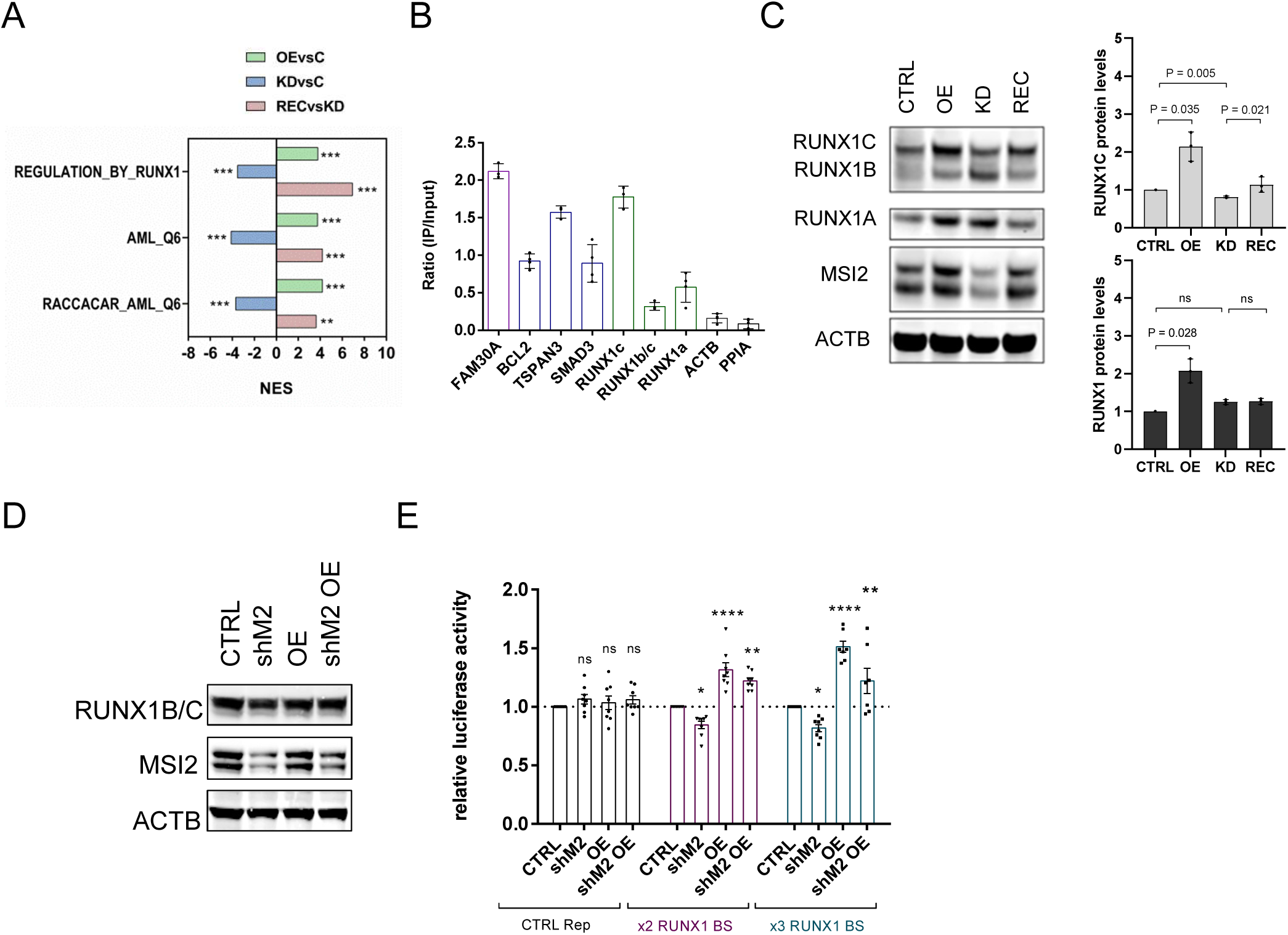
The *FAM30A*/MSI2 axis regulates RUNX1 transcriptional activation. **A.** GSEA analysis (C2, MSigDB) in KG-1a stable cell lines shows a significant enrichment for RUNX1 activation. **B.** Immunoprecipitation of MSI2 in KG-1a cells and subsequent RT-qPCR analysis of MSI2-associated RNAs, depicting MSI2-targets (blue), RUNX1 isoforms (green) and negative controls (black). **C.** Representative Western blot (left) for MSI2 and RUNX1 isoforms in stable KG-1a cell lines and quantitative analysis (right) of RUNX1C and total RUNX1 protein levels, normalised to control cells. ACTB was used as a loading control. **D.** Western blot analysis in stable K562 cell lines for RUNX1 and MSI2 protein. **E.** Quantitative analysis of relative luciferase activity in stable K562 cell lines transfected with constructs comprising RUNX1 binding sites to assess RUNX1 activation. Normalization was performed against control cells and the control reporter (CTRL Rep). Statistical significance was calculated using unpaired two-tailed student t-test (* P < 0.05, ** P < 0.01, *** P < 0.001, **** P < 0.0001, ns – not significant). Data is displayed as mean values and error bars represent SEM.

### *FAM30A* KD abrogates murine BM engraftment, partially restored by *FAM30A* repeats

To evaluate how *FAM30A* and its repeats regulate engraftment or the ability to initiate and maintain leukaemia in mice, we cultured the KG-1a stable cell clones and transplanted them into 8-12 weeks sub-lethally irradiated NRG-SGM3 transgenic mice (3 x 10^6^ cells per mouse, average n=6 per group) (Figure 6A). To provide evidence of leukaemia-like disease, cells from murine bone marrow (BM) and spleen were isolated to determine the levels of human cells (Figure 6B). In addition, spleen weights were also quantified (Supplementary Figure 6A). Mice transplanted with control cells successfully engrafted in the BM (42.5±10.3%) and spleen (26.9±11.9%). Importantly, *FAM30A* KD abrogated murine engraftment compared to control (BM: 0.5±0.2%, Spleen: 0.5±0.1%) after 6 weeks, which was partially restored in the REC condition (BM: 6.8±4.1%, Spleen: 1.1±0.7%) when *FAM30A* repeats were ectopically expressed in the KD setting (Figure 6C, D). Transplanted cells overexpressing the *FAM30A* repeats led to significantly lower murine engraftment in mice (BM: 14.8±4.1%, Spleen: 1.2±0.5%) compared to controls. To explore the reduced engraftment potential in the OE setting, we analysed endogenous *FAM30A* levels in the stable cell lines used for the xenograft experiments at different time points in culture. RT-qPCR showed that endogenous *FAM30A* was significantly downregulated when *FAM30A* repeats were overexpressed in long term cultures (Figure 6E), which likely accounts for the lower engraftment levels in mice as well. Connected to this, CD34/CD11b-staining in these *in vivo* engrafted cells suggested that *FAM30A* repeat overexpressing cells have initiated myeloid differentiation compared to control (Figure 6F). Our engraftment studies provide evidence that *FAM30A* repeats alone can only partially recapitulate the effects seen upon depletion of the full *FAM30A* lncRNA. Taken together, these results further emphasize the critical role of *FAM30A* in governing LSC features like murine BM and spleen engraftment, a defining feature of fully competent LSCs in AML.

**Figure 6.**
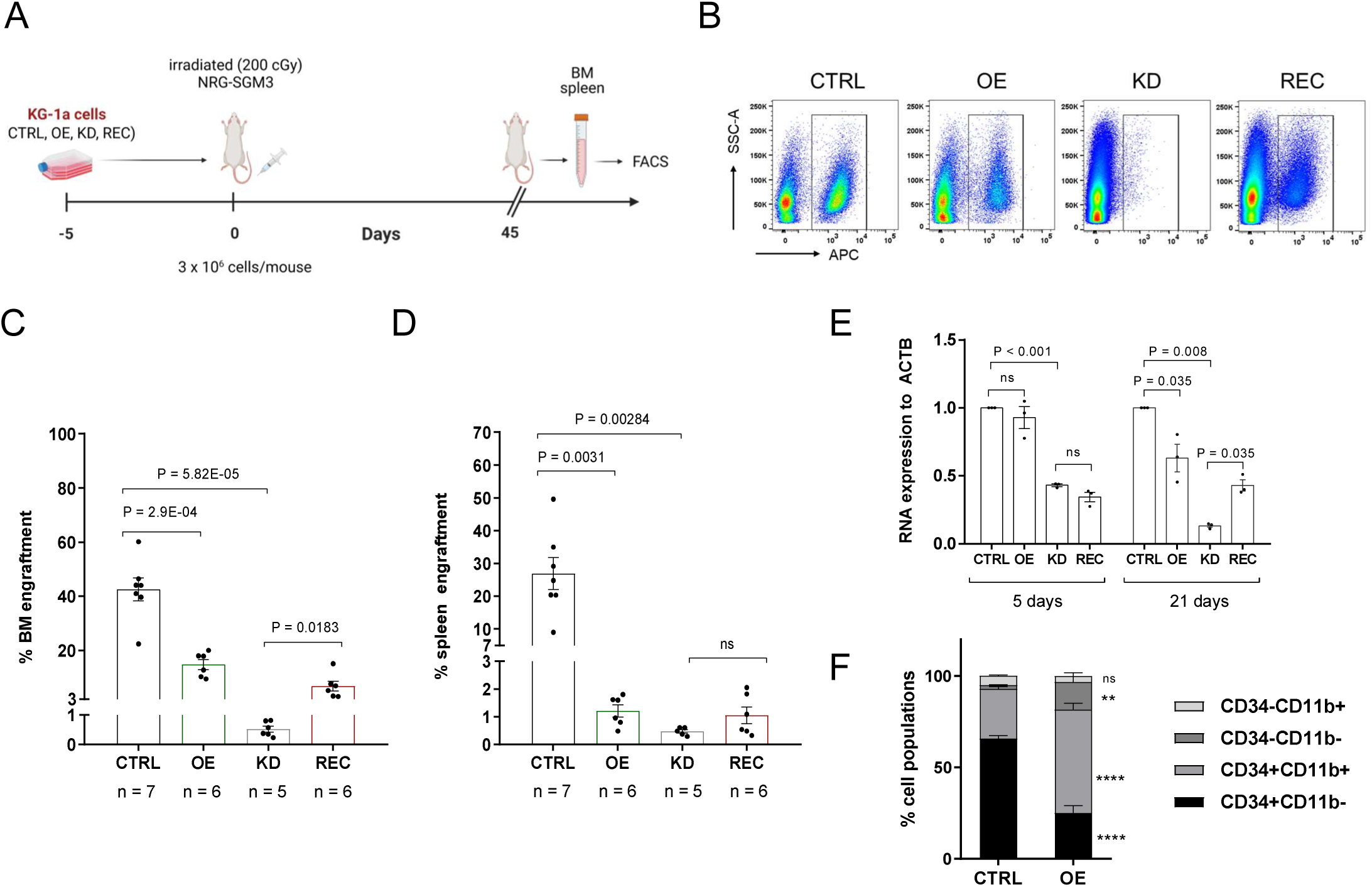
*FAM30A* depletion abrogates murine BM engraftment after 6 weeks post-transplant. **A.** Schematic representation of the study design testing murine BM and spleen engraftment upon modulation of *FAM30A* expression in immunodeficient mice. **B.** Representative FACS analysis for the assessment of engraftment of the stable KG-1a cell lines as percentages of APC-positive cells. **C.** Quantification of the percentage of APC-positive cells in murine BM. **D.** Quantification of the percentage of APC-positive cells in murine spleen. **E.** RT-qPCR analysis for *FAM30A* levels in stable KG-1a cell lines grown in liquid culture at two different time points resembling before/after transplant, normalized to control cells. **F.** Murine BM samples were stained for human CD34 and CD11b after transplantation in respective cohorts (CTRL, OE) and analysed by FACS. Statistical significance was calculated using unpaired two tailed student t-test (** P < 0.01, **** P < 0.0001, ns – not significant). Data is displayed as mean values and error bars represent SEM.

## DISCUSSION

Deregulated lncRNA expression is increasingly been acknowledged as a key factor in AML pathogenesis^10, 45^. The herein presented study provides the first functional characterization of the stem cell-specific lncRNA *FAM30A* in AML cells. By employing an AML LSC-like cell line model with high *FAM30A* expression, *in vitro* and *in vivo* experiments revealed that modulation of *FAM30A* and its newly identified *FAM30A* repeats have a direct effect on clonogenicity, chemoresistance and engraftment of AML cells. Mechanistically, we also revealed that *FAM30A* repeats associate and regulate MSI2 protein levels and its downstream target SMAD3, important for LSC maintenance and leukaemia progression^42^. We also propose that *FAM30A* repeats drive higher MSI2-association especially to the RUNX1C isoform, which is likely to facilitate overall increased RUNX1 transcriptional activation.

Located at the 14q32.33, the *FAM30A* transcripts are embedded in antisense orientation to the variable region of the immunoglobulin heavy chain (*IGHV*) locus, necessary for the production of antibodies, which could explain the relatively high expression levels of *FAM30A* in B-cells^27^ and its prevalence in B-cell leukaemia (Figure 1A). Interestingly, chromosomal rearrangements involving this genomic region result in a predisposition towards the onset of myeloid immature acute leukaemia^46^. These findings might explain the higher *FAM30A* levels in the more primitive CD34+CD38-LSC fraction which is characterized by higher self-renewal and chemoresistance properties^16, 28^. Nevertheless, the relatively high *FAM30A* levels detected in early HSCs and myeloid progenitors (Figure 1G) imply that *FAM30A* could be relevant for both HSCs and LSCs, as shown for other AML-relevant lncRNAs in previous studies^12, 13^.

Repeated RNA sequences in lncRNAs have been shown to represent functional RNA elements as many of these lncRNAs depend on recruitment of several RBPs (i.e. MSI2) to exert their function^47, 48^. The six tandem repeats of *FAM30A* represent an element of *FAM30A* that already caught attention decades ago^49^, although there was no reported function so far. Conservation studies performed in this study suggest a selective evolutionary pressure to increase the number of a single *FAM30A* repeat over time. Since one *FAM30A* repeat comprises three putative MSI2 binding sites, this suggests that one *FAM30A* molecule holds the capacity to potentially bind a multitude of MSI2 molecules simultaneously. This underscores the biological relevance of *FAM30A* and MSI2 not only in the LSC context but also potentially in normal haematopoiesis. The present study also provides the first evidence that *FAM30A* is a cytoplasmic lncRNA lacking polysome association. Nevertheless, our results cannot rule out that ribosomes transiently bind to *FAM30A*, as a substantial proportion of *FAM30A* (≈25%) was observed in complexes co-migrating with monosomal fractions. Preliminary *in silico* prediction of lncRNA-encoded peptides (LncPep tool) detected putative micropeptides ranging from 50-150 amino acids (data not shown) that could contribute to *FAM30A*-mediated phenotypes. This hypothesis has been substantiated in a recent study that showed that an lncRNA-encoded micropeptide named *APPLE* promotes an oncogenic translation program by promoting association of RBPs^50^.

The relevance and requirement of MSI2 as a key regulator in HSCs and LSCs in AML is well established^41, 42, 51, 52^. To our knowledge, there have been no reports to date of an lncRNA that directly associates with and positively influences MSI2 protein function in leukaemia. Only *LINC00942* has been identified as an lncRNA that binds and regulates MSI2 protein degradation to promote chemoresistance in gastric cancer^53^. *FAM30A* KD correlated with downregulation of canonical AML LSC signatures, decreased cell viability, increased apoptosis and sensitivity to chemotherapeutics compared to control. Importantly, *FAM30A* depletion abrogated BM engraftment by 6 weeks post-transplant suggesting that it serves as a key regulator of engraftment potential. These phenotypic changes were partially rescued by exogenous expression of the MSI2-interacting region of *FAM30A.* These results suggest a collaborative complex involving *FAM30A* and MSI2 within the stem cell compartment, as analysis of scRNAseq data also suggested (Figure 1G, Supplementary Figure 4B). We hypothesize that this *FAM30A*/MSI2 complex follows previous results performed in cancer stem cells showing how an lncRNA can directly affect protein levels of their cognate RBP and affect stability of its target mRNAs^54, 55^.

Interestingly, a recent study that used scRNAseq to analyse the heterogeneity of LSC transcriptional reprogramming after chemotherapy in primary refractory and relapsed AML patients found that both *FAM30A* and MSI2 were among the most highly upregulated genes. These genes were particularly elevated in a quiescent subpopulation of LSCs responsible for chemoresistance across different AML subtypes^36^. The effects seen in our study after depletion and/or overexpression of *FAM30A* repeats aligns with previous results that showed how MSI2 decreased/enhanced activity affects self-renewal, chemoresistance, colony-formation potential, apoptosis, myeloid differentiation and engraftment in AML cells^41, 52, 56–58^. Moreover, GSEA analysis showed that some pro-LSC signalling pathways controlled by MSI2 (i.e. TGF-β/SMAD3^52^) were positively correlated with changes in levels of *FAM30A* repeats. Nevertheless, we also cannot exclude the possibility that *FAM30A*/MSI2 require the association of additional factors to exert their function. Indeed, LC/MS-MS analysis following RNA pulldown of *FAM30A* repeats identified other RBPs of potential relevance to AML LSC function, such as SYNCRIP^57^.

The transcription factor RUNX1 is essential in HSCs^59^ and AML progression^60^. From the RUNX1 gene, the three main isoforms (A,B,C) are produced through alternative splicing as well as alternative promoter usage and all of them have been shown to contribute to leukaemia progression at different stages and subtypes^61, 62^. The RUNX1C isoform has attracted considerable attention as the stem cell-specific isoform in AML for its relevant role in the maintenance of CD34+ LSCs (i.e. frequency and engraftment)^43^. Moreover, RUNX1C has been shown to be important in the myeloid compartment as its enforced expression promotes colony-formation potential in myeloid progenitors (i.e. CFU-GM, CFU-M)^63^. In our study, we reveal not only that MSI2 strongly binds to RUNX1C but we also show that increased RUNX1C protein levels and RUNX1 transcriptional activity can be mediated by both *FAM30A* and MSI2. This suggests that the phenotypic changes in LSC features resulting from overexpression of the MSI2-interacting region of *FAM30A* may be due to elevated RUNX1 levels and activity. A computational study identified *FAM30A* as a mediator gene influencing the survival of AML patients with RUNX1 alterations suggesting a potential self-regulatory feedback loop^64^. In addition to the RUNX1 transcriptional pathway other signalling networks could play important functions in contributing to *FAM30A*’s cellular roles. This could include the IL2-STAT5 pathway or the TNFA-pathway, since they were affected by *FAM30A* modulation and have been reported to influence AML pathogenesis as well (Figure 3D)^65, 66^.

Notably, although overexpression of *FAM30A* repeats enhanced pro-LSC features *in vitro* and in the short term, it led to reduced murine BM engraftment potential compared to the control. This reduction may be linked to lower endogenous *FAM30A* levels in the long-term and more differentiated AML-like blasts.

In conclusion, our data supports an extensive role for the lncRNA *FAM30A* in LSC biology of AML cells. We propose that it is mechanistically driven by the interaction of *FAM30A* repeats with MSI2 which affects also RUNX1 activation, thus implying a novel positive feedback loop relevant for LSC maintenance in AML cells. Interestingly, a small-molecule inhibitor targeting MSI2 *in vivo* has been shown to reduce disease burden in murine AML model while sparing normal CD34+ cord-blood cells thus this opens a therapeutic window where *FAM30A* and MSI2 association can be selectively targeted without affecting normal haematopoiesis^58^. We propose that a further exploration of this mechanism is warranted as a means of defining a therapeutically tractable pathway for eradicating LSCs, which could significantly improve therapy response in AML patients.

## MATERIALS and METHODS

### Cell lines and cell culture

The origin of the cell lines is summarized in Supplementary Table S1. GDM-1, KG-1a, K562, ME-1, MOLM-13 and MV4-11 were cultured in Roswell Park Memorial Institute medium (RPMI-1640, Thermo Fisher Scientific) supplemented with 20% fetal bovine serum (FBS), 1% GlutaMAX (L-Alanyl-L-glutamin) and 1% (v/v) penicillin-streptomycin. At every passaging, fast-growing cells (MV4-11, MOLM-13 ME-1, K562) were usually seeded at lower cell density (0.1-0.5 x 10^6^ cells/ml) and slow-growing and more stem-like cells (GDM-1 and KG-1a) were maintained at higher cell density (0.5-1 x 10^6^ cells/ml). For the adherently growing HEK293T17 cell line, the culture conditions were in Dulbecco’s modified Eagle’s medium (DMEM, Thermo Fisher Scientific) supplemented in 10% FBS and 1% GlutaMAX. Prior to passaging, the medium was discarded and cells were washed with 1x PBS. Following this, cells were detached witch 0.05% Trypsin/0.4mM EDTA and the process was stopped by the addition of fresh DMEM supplemented with 10% FBS. All cell lines were maintained in an incubator at 37°C and 5% CO_2_.

### Lentiviral transduction

For lentivirus production, HEK293T17 cells were co-transfected with the packaging plasmids psPax2 (Addgene), pMD2.G (Addgene) and indicated lentiviral pLVX vectors (Supplementary Table S2). Cells expressing both RFP (pLVX_RFP-plasmids) and Crimson (pLVX_Crimson_shRNA-plasmids) were sorted using FACS Melody (BD Biosciences) cell sorter. Lentiviral supernatant was collected 24–48 h post-transfection. Viral titers were determined 72 h post-infection using HEK293T17 cells by flow cytometry (RFP or Crimson) and a MACS Quant Analyzer (Miltenyi BioTech). For lentiviral transduction cells were infected at 10 MOI (multiplicity of infection) for 72 h.

### Subcellular fractionation

For fractionation, 1.5LJxLJ10^6^ cells per condition were harvested in fractionation buffer (10 mM Hepes-KOH pH 7.2, 150 mM KCl, 5LJmM MgCl_2_) supplemented with 200 μg/ml digitonin (Sigma-Aldrich, St. Louis, MO, USA) for KG-1a cells. Subcellular fractions were separated by centrifugation at 1000 g for 2 min. Supernatants containing the cytoplasmic fraction were transferred to a new tube. The remaining pellet was washed with fractionation buffer supplemented with digitonin and centrifuged at 1000 g for 2 min. Pellets containing the nuclear fraction were lysed in total lysis buffer (50 mM Tris–HCl pH 7.4, 50 mM NaCl, 1% SDS, 1LJmM MgCl_2_, Turbo nuclease [Jena Bioscience, 250 U/μl]). Subcellular fractionation was confirmed by detecting EEF2 in the cytoplasmic and PTBP1 in the nuclear fraction.

### Sucrose-based gradients and targeted polysome profiling

For the isolation of polysome fractions and profiling, wild-type KG-1a cells were cultured at a concentration of 0.5-1 x 10LJ cells/ml. The day before, a sucrose gradient (15-45%) diluted in gradient buffer (10 mM HEPES-KOH pH 7.2, 5 mM MgCl_2_, 150 mM KCl) was carefully layered into ultracentrifugation tubes. This involved preparing five sucrose solutions at concentrations of 15%, 22.5%, 30%, 37.5%, and 45%. Starting with the 45% solution, 2 ml of each subsequent solution was layered sequentially, creating a total volume of 10 ml. The tubes were then left overnight at 4°C to allow the gradient to diffuse properly. The following day, cells were harvested, washed with PBS, lysed in lysis buffer (gradient buffer with 0.5% NP-40) for 5 min, and centrifuged at 13,000 rpm for 5 min in a tabletop micro-centrifuge. The supernatant was collected and kept on ice for further polysome profiling. An equivalent of 10 million previously lysed KG-1a cells was loaded onto 11 ml of the 15-45% linear sucrose gradient. The gradients were centrifuged at 40,000 rpm for 2 h at 4°C using a SW40 Ti rotor (Beckman Coulter). Eleven fractions of 500 µl each were collected and 200 µl and 50 µl were used for RNA and protein analysis, respectively. For RNA analyses, 0.2 ml of each fraction was mixed with 1 ml of TRIzol and processed using standard RNA extraction protocols. Unfractionated cell lysates were used as input to assess transcript abundances across each fraction of the gradient.

### Biotinylated RNA-pulldown and quantitative proteomics by LC-MS/MS

To conduct RNA pulldowns, 15 μl of Dynabeads™ MyOne-Streptavidin-T1 (Thermo Fisher Scientific) per pulldown were placed into a reaction tube. The beads were washed three times with an equal volume of BW buffer (5 mM Tris-HCl pH 7.5, 0.5 mM EDTA, 1 mM NaCl) using a magnetic rack and then incubated with 0.5-1 μg of *in vitro* transcribed RNA. As a control, beads were incubated with no RNA for the beads control (namely BC) for 20 min at room temperature. Concurrently, cells were washed with PBS and lysed (gradient buffer with 0.5% NP-40 for Western blot or 1% DDM for MS analyses) for 10 min and centrifuged for 5 min at 13,000 rpm. The resulting supernatant was transferred to a new reaction tube, and 20 μl (1:10) of the lysate was used for the input sample quantitative proteomics (LC-MS/MS analysis). Meanwhile, 200 μl of the cell lysate was mixed with each of the RNA-coupled streptavidin beads and incubated for 45 min at room temperature. Following this, pulldown samples were washed three times with 200 μl of gradient Lysis buffer and separated on a magnetic rack to eliminate nonspecific proteins and RNA. For the analysis of associated proteins, the beads were re-suspended in 50 μl of 1x NuPage™ LDS sample buffer (Thermo Fisher Scientific) diluted with lysis buffer and boiled at 95 °C for 5 min to elute proteins. Then, samples were separated from the beads on a magnetic rack and subsequently analysed for downstream protein analysis through SDS-PAGE and Western blot.

Pulldown samples (three replicates) were prepared for liquid chromatography tandem mass spectrometry (LC-MS/MS) following a modified FASP (filter-aided sample preparation) protocol^67^. Briefly, protein samples were incubated and washed with 8 M urea in 50 mM HEPES (4-(2-hydroxyethyl)-1-piperazineethanesulfonic acid), 10 mM TCEP, pH 8.5. Washing steps were performed using 0.5-ml centrifugal filter units (30-kDa cut-off, Sigma Aldrich, Darmstadt, Germany) and two centrifugation steps (14,000g, 10 min). Afterwards, alkylation of proteins was performed with 50 mM iodoacetamide in 8 M urea, 50 mM HEPES, pH 8.5 at room temperature (20 min in the dark). Samples were washed twice with 50 mM HEPES, pH 8.5 (18,000g, 2 x 10min) before they were incubated with 1 μg trypsin (Promega) in 50 mM HEPES, pH 8.5 at 37°C overnight. After enzymatic digestion, samples were acidified with trifluoroacetic acid (TFA) at 0.5% (v/v) final concentration.

Peptide solutions were analysed by (LC-MS/MS) on an Ultimate 3000 RSLC nano-HPLC system coupled to a Q-Exactive Plus mass spectrometer with a Nanospray Flex ion source (all from Thermo Fisher Scientific). Samples were loaded onto an RP C18 pre-column (Acclaim PepMap, 300 μm x 5 mm, 5 μm, 100 Å, Thermo Fisher Scientific) and washed with 0.1% (*v/v*) TFA for 15 min at 30 μl/min. Peptides were eluted from the pre-column and separated on a 200-cm μPAC C18 separation column (Pharmafluidics) that had been equilibrated with 3% solvent B (solvent A: 0.1% (*v/v*) FA (formic acid), solvent B: ACN, 0.1% (*v*/*v*) FA). A linear gradient from 3–35% solvent B within 180 min at 600 nL/min (0–30 min) and 300nl/min (30–180 min) was used to elute peptides from the separation column. Data were acquired in data-dependent MS/MS mode, HCD (higher-energy collision-induced dissociation) with normalised collision energy (NCE) of 28% was used for fragmentation. Each high-resolution full MS scan (*m*/*z* 375 to 1799, *R* = 140 000, target value (TV) 3 000 000, max. injection time (IT) 50 ms) was followed by high-resolution (*R* = 17500, TV 200 000, IT 120 ms, isolation window 2Th) product ion scans, starting with the most intense signal in the full scan MS, dynamic exclusion (duration 30 s, window ± 3ppm) was enabled. For peptide identification and quantification, data were searched against the Swiss-Prot database (tax. *Homo sapiens*, version 11/19; 20 315 entries) using Proteome Discoverer, version 2.4 (Thermo Fisher Scientific). A maximum mass deviation of 10 ppm was applied for precursor ions, while for product ions, max. 0.02 Da were allowed. Oxidation of Met and acetylation of protein N-termini were set as variable modifications. Carbamidomethylation of cysteines was included as fixed modification. A maximum of two missed cleavage sites were considered for peptides. Peptides were quantified via LFQ (label-free quantitation) and data were finally subjected to quantile normalisation to derive protein enrichment for *FAM30A* repeats.

### RNA immunoprecipitation

KG-1a cells (8 x 10^6^ per condition) were harvested and washed with PBS and then lysed using lysis buffer for immunoprecipitation (10 mM HEPES-KOH pH 7.2, 5 mM MgCl_2_, 150 mM KCl and 1% Triton X-100). Inputs for either RNA or protein were extracted from the lysates. Then, magnetic Protein G Dynabeads (Life Technologies) were washed three times with Immunoprecipitation buffer and 5-10 μg/condition of anti-MSI2 (Bethyl) was added and incubated for 15 min. The resulting supernatant with the protein lysate was incubated with the antibody-bound beads for 60 min at room temperature on a spinning-wheel. Following three washing steps with Immunoprecipitation buffer, protein–RNA complexes were eluted using 100 µl of Immunoprecipitation buffer supplemented with 1% SDS and incubated at 65°C for 10 min and beads were separated in the magnetic rack. The supernatant containing the eluted protein-RNA complexes was transferred to a new tube and 20 µl was used for protein analysis and 80 µl for RNA analysis. Protein enrichment was subsequently analysed by Western blot and co-purified RNAs were extracted using TRIzol and subjected to analysis by RT-qPCR.

### Cell viability, hypoxia and colony-formation assays

Cell viability was assessed using the CellTiter-Glo® Luminescence Cell Viability Assay (Promega, Madison, WI, USA) following the manufacturer’s instructions. Cells were seeded at a concentration of 0.5-2 x 10^5^ cells/ml in a 24-well plate and cell viability was determined after 48-72 hours. Then, CellTiter-Glo® reagent was added to the cell culture medium with cells at a 1:1 ratio and incubated for 10 min. The resulting luminescence signal generated by the luciferase reaction, was detected using a GloMAX® luminometer (Promega).

Prior seeding cells in hypoxia conditions, cells were washed with PBS and seeded at a cell density of 0.5 x 10^6^ cells/ml. Then, cells underwent hypoxic treatment (1% O_2_, 5% CO_2_) which was performed in a humidified incubator (Innova CO-170; New Brunswick Scientific) and cells were maintained for 72 h. Afterwards, cell counts and the percentage of living cells were determined using a cell counter.

For colony-formation assays, cells were seeded at 5000 cells/ml in 3 ml MethoCult^TM^ (Stem Cell Technologies, H4034) in a 3.5 cm culture dish and placed in an incubator at 37 °C and 5% CO_2_. Before seeding, the clonogenic potential (CFC-outputs) was calculated based on cell expansion in liquid culture for 48 h. After 14 d of incubation, the number of colonies was manually counted and classified into each colony type following manufactureŕs instructions.

### Apoptosis assay

To measure apoptotic levels after treatment with chemotherapeutics, cells were seeded and incubated for 48 h before analysis. Afterwards, cells were pelleted and washed twice with PBS and incubated with the staining agents Annexin-V-FITC (1:2000) and DAPI in a solution of PBS with 1% Bovine Serum Albumine (BSA) for 1h at 4°C in the dark. Annexin-V-positive cells were used as early apoptosis and Annexin-V-positive, DAPI-positive cells as late apoptosis indicators, respectively. Then, cells were washed again with PBS and plated in a 96-well plate for subsequent analysis of early and late apoptosis using flow cytometry with a MACSQuant flow cytometer and the MACSQuantify Software (Miltenyi Biotec).

### Luciferase reporter assays

K562 stable cell lines were seeded one day before transfection at a cell density of 2.5 x 10^5^ cells/ml in a 24-well plate. After one day, cells were transiently transfected using DharmaFECT kb with pmirGLO Promo+Stopp*HindIII plasmid (empty vector) comprising two and three palindromic RUNX1 binding sites (TGTGG) in the promoter region of the Firefly luciferase gene (see Supplementary Table S2). Renilla luciferase on the same plasmid served as the normalization control. The luminescence values from Firefly and Renilla luciferases were determined 48 hours post transfection by DualGLO (Promega) according to the manufacturer’s protocol.

### RNA isolation and quantitative RT-PCR

RNA extraction was carried out with TRIzol (Thermo Fisher Scientific) from cell pellets that were mechanically disrupted before RNA isolation. For cDNA synthesis, 2 μg of total RNA was used as a template, with random primers and the M-MLV reverse transcription system (Promega). Quantitative PCR was then conducted using SYBRgreen I technology with ORA qPCR Green ROX L Mix (HighQu, Kraichtal, Germany) on a LightCycler 480 II 384-well system (Roche, Basel, Switzerland). RNA levels were measured using the ΔΔCt method, normalized to ACTB and EEF2, with primers listed in Supplementary Table S4.

### RNA-sequencing and data processing

For bulk RNA-sequencing (RNA-seq) 500 ng of total RNA of stable KG-1a cell clones were used (n=3 in all cell samples). Library preparation and sequencing were performed by Novogene (Hong Kong) on an Illumina HiSeq platform. First, low-quality read ends as well as remaining parts of sequencing adapters were clipped using cutadapt (v1.4; 2.8). Subsequently, reads were aligned to the human genome (UCSC GRCh38/hg38) using hisat2 v2.1.0^68^. Featurecounts (v1.5.3; 2.0)^69^ was used for summarizing gene-mapped reads. Ensembl (GRCh38.89; GRCh38.102)^70^ was used for annotations. Differential gene expression (DE) was determined by the R package edger (v3.28.0; 3.34)^71^ using TMM (trimmed mean of M values) normalization on raw count data. RNA expression values were obtained as FPKM (fragments per kilobase million mapped reads) values.

### Protein extraction and Western blot

For total protein extraction, cell pellets were lysed in total lysis buffer (50 mM Tris–HCl pH 7.4, 50 mM NaCl, 1% SDS, 1LJmM MgCl_2_, Turbo nuclease [Jena Bioscience, Jena, Germany, 250 U/μl]). The protein concentration of each sample was determined using a colorimetric assay (Bio-Rad, Hercules, CA, USA) according to manufacturer’s protocol. Equal amounts of total protein (30 μg) were separated by NuPAGE Bis-Tris (4–12%) gels (Thermo Fisher Scientific) and transferred to a nitrocellulose membrane (Amersham, GE Healthcare, Chicago, IL, USA) using the Mini Gel Tank Blotting system (Thermo Fisher Scientific). The membranes were blocked for 1h at room temperature in phosphate-buffered saline solution (PBS; 140 mM NaCl, 2.6 mM KCl, 1.5LJmM KH_2_PO_4_ and 8LJmM Na_2_HPO_4_) containing 5% (w/v) skimmed milk. Membranes were incubated with primary antibody at 4 °C overnight, washed with PBS supplemented with 0.1% Tween-20, incubated with secondary antibody at room temperature for 1 h and analysed using the Odyssey infrared scanner (LI-COR, Lincoln, NE, USA). Primary and secondary antibodies are listed in Supplementary Table S6.

### Conservation analysis

To assess the phylogenetic conservation of *FAM30A* repeats across vertebrates, the sequence of a single *FAM30A* repeat (76 and 78 base pairs) was downloaded from NCBI gene website (https://www.ncbi.nlm.nih.gov/gene/) using its annotated sequence (NR_026800.2) and subsequently blasted against all taxonomic nucleotide databases using NCBI nblast tool (https://blast.ncbi.nlm.nih.gov/). Following the alignment of a single repeat through the NCBI nblast tool against various taxonomic nucleotide databases, the sequence exhibited alignment primarily with analogous sequences found in 21 primate species. Thus, the genome sequences of these species were downloaded from NCBI and ENSEMBL (https://www.ensembl.org). Then, in order to analyse the context of the repeat, longer sequences including the flanking regions of the repeats were downloaded (500 base pairs upstream and downstream) and blasted against the primate genomes where the *FAM30A* repeat was detected. After multiple sequence alignment using M-Coffee software^72^ a conservation score was extracted from 0 (lowest) to 9 (highest) for each nucleotide. The conservation along the 1500 base pairs was represented in a curve after locally weighted regression by Lowess algorithm using GraphPad 8.0.

### Xenotransplantation of KG-1a cells, animal handling and ethics approval

Human cell xenograft experiments were performed under complete supervision of Prof. Dr. David Vetrie (Wolfson Wohl Cancer Research Centre, University of Glasgow, United Kingdom) and his laboratory staff. All mice were housed at the Beatson Research Unit (License holder: Karen Keeshan, PPL number: PP4496278, University of Glasgow, UK) and experimental protocols were approved by Karen Keeshan (Wolfson Wohl Cancer Research Centre, University of Glasgow, United Kingdom). Before engraftment, KG-1a stable cell clones were cultured in RPMI with 20% FBS for five days for optimum cell growth at 37°C and 5% CO_2_. KG-1a cells (3 × 10^6^ cells per mouse) were diluted in ice-cold PBS and transplanted via tail vein injection into 8-12-week-old sub-lethally irradiated NRG-SGM3 immune-compromised mice (NOD.Cg-Rag1^tm1Mom^ Il2rg^tm1Wjl^ Tg(CMV-IL3,CSF2,KITLG)1Eav/J). After 6 weeks upon transplantation, all mice were euthanized, and bone marrow (BM) was harvested from the hind legs (ilia, femurs, and tibias). BM was filtered and analysed by flow cytometry using human anti-CD34-BV510 and human anti-CD38-VioBlue (see table 5). Engrafted human cell immunophenotypes were determined by flow cytometry (BD, FACSVerse). To assess level of engraftment of human cells, cells were gated for APC-positive cells as KG-1a stable clones emit RFP/Crimson fluorescence and also using transplanted mice as negative control (with no fluorescence).

### Public data, software and databases

MSI2 (Musashi-2) CLIP peak data of K562 and NB4 cells were obtained from the Gene Expression Omnibus (GEO accession numbers: GSE62115; GSM2448651; GSE69583). Peaks from the obtained 6 experiments were annotated to gene features (5’ UTR, CDS and 3’ UTR), summarized and visualized via an in-house R-script.

Western blot images were visualized, and the band intensities calculated using Image Studio™ Lite Quantification Software (LI-COR Biosciences). In case the orientation, brightness or contrast of the images needed adjustment, Adobe Photoshop or GIMP software were used. To visualize the sequencing files provided by Eurofins and to create plasmid maps, SnapGene software was used. The flow cytometry results were analysed and graphs depicted using FlowJo™ v10.8 software (BD Life Sciences). For Figure 2A, 6A and Supplementary Figure 2D BioRender.com was used. RNA sequencing was visualised with the Integrative Genomics Viewer (IGV).

### Gene set enrichment analysis and functional annotation clustering

The Gene Set Enrichment Analyses (GSEA) were conducted using GSEA software (V3.0)^73^ on pre-ranked lists, with a focus on Hallmark and Gene Ontology (Biology Processes) gene sets from MSigDB (v6.2) as well as AML and LSC signatures. The ranking of all protein-coding genes was based on fold changes observed in OE vs CTRL, KD vs CTRL and REC vs KD as determined by RNA-seq. A permutation number of 1000 was implemented, and classical enrichment statistics were chosen for the analysis.

The potential proteins bound by *FAM30A* repeats RNA versus beads control (no RNA) were clustered based on identified peptides (fold-change, P < 0.05) derived from LC-MS/MS analysis and uploaded onto the Database for Annotation, Visualization, and Integrated Discovery (DAVID) (http://david.abcc.ncifcrf.gov/). From the proteins significantly linked with *FAM30A* repeats RNA, a gene list was derived and underwent Gene Ontology (Biology Process) classification and the most significant biological processed were selected and depicted.

### Statistical analyses

All experiments were conducted using a minimum of three independent biological replicates, unless otherwise stated. Statistical analyses were conducted using GraphPad Prism software (V8.0) or Microsoft Excel. Statistical significance was determined using unpaired two-tailed Student’s t-test, unless specified. Statistical analysis for phylogenetic conservation of the *FAM30A* repeats across vertebrates was performed using locally weighted regression of curve by Lowess algorithm in GraphPad Prism. Gene expression correlations were assessed via Pearson correlation using the cBioPortal for Cancer Genomics website (cbioportal.org), or the Logrank test (Mantel– Cox) in GEPIA2 (gepia2.cancer-pku.cn) and Kaplan Meier Plotter (kmplot.com).

## ACKNOWLEDGMENTS and FUNDING

This work was supported by grants from the Deutsche Forschungsgemeinschaft [DFG, German Research Foundation; RTG 2467, project number 391498659 ‘Intrinsically disordered proteins—molecular principles, cellular functions, and diseases’; FOR 5433, project number 468534282 ‘RNA in focus (RIF): from mechanisms to novel therapeutic strategies in cancer treatment’] and the Wilhelm-Roux-Program (Medical Faculty, Martin-Luther-University Halle–Wittenberg).

The authors would like to thank the team of the Core Facility Imaging (CFI, Medical faculty, University of Halle-Wittenberg) for broad support with image acquisition and data analysis.

## SUPPLEMENTARY FIGURE LEGENDS

**Supplementary Figure 1. Mutational spectrum and subtype-specific pattern of *FAM30A* expression in AML patients. A.** Normalised *FAM30A* (ENSG00000226777) expression levels in a variety of tumour samples extracted TCGA database. **B.** *FAM30A* expression data extracted from TCGA (PanCancerAtlas) assigned to each of the karyotype based on patient data. **C.** *FAM30A* levels derived from R2 platform (Bohlander, n=422) with AML subtypes assigned according to the FAB classification. **D.** RNA levels of *FAM30A* among AML patients harbouring the most common mutations extracted from patients data as in C. **E.** UMAP visualization map of *FAM30A* and other well-known HSC/LSC markers (CD34, HOPX) clustered based on cell-types from scRNAseq portal (Broad Institute)^38^. Statistical significance was calculated using unpaired two-tailed student t-test (** P < 0.01, **** P < 0.0001, ns – not significant). Data is displayed as mean values and error bars represent SEM. + represents mutations in the gene, wt = wild-type; ITD = internal tandem duplication; TKD = mutation in the tyrosine kinase domain.

**Supplementary Figure 2. Identification *FAM30A* repeat region. A.** Taxonomic tree of primates showing the genome sequences of the species used for each clade. Light blue highlights species where at least one *FAM30A* repeat was detected, while dark blue marks species with more than one *FAM30A* repeat. **B.** Quantitative analysis of the conservation score extracted from multiple sequence alignment (M-Coffee) of a sequence range (500 bp upstream/downstream the repeat region). For comparison, five primate genomes were extracted (NCBI, Ensembl). Statistical analysis was performed using locally weighted regression of curve by Lowess algorithm. Highlighted in red is the beginning of the Repeat 1 in the *FAM30A* sequence. **C.** Normalised *FAM30A* (ENSG00000226777) expression levels across a variety of normal tissues extracted from GTEX. **D.** Illustrative depiction of the oligonucleotides used for detection of *FAM30A* (higher panel), quantification via RT-qPCR analysis (lower left panel) and depiction from IGV software of RNA-sequencing performed in KG-1a cells. The image shows the main five exons of *FAM30A* with the highest raw reads.

**Supplementary Figure 3. Modulation of *FAM30A* levels affects resistance to hypoxia and treatment with chemotherapeutic drugs. A.** Western blot analysis of stable KG-1a cell lines after being cultured in normoxic (21% O_2_) and hypoxic (1% O_2_) culture conditions. HIF1a was used as hypoxia marker and Lamin B1 as loading control under normoxia and hypoxia, respectively. Normalisation was performed by comparing to control cells under hypoxia. **B.** Representative FACS analysis of stable KG-1a cell lines stained with Annexin-V/DAPI after daunorubicin treatment (0.1 µM, 48 h). **C.** Quantification of the percentage of cells in early and late apoptosis from B. **D.** Quantitative analysis of the cell viability of stable MV4-11 and ME-1 OE cells treated with cytarabine (3 µM) and daunorubicin (0.1 µM) after 48 h. Normalization was performed against control cells. **E.** Representative FACS analysis of CD11b-positively stained stable KG-1a cell lines after 5 days in liquid culture. Statistical significance was calculated using unpaired two-tailed student t-test. Data is displayed as mean values and error bars represent SEM.

**Supplementary Figure 4. *FAM30A* expression highly correlates with MSI2 and other stem cell markers. A.** RNA levels of *FAM30A* and MSI2 in AML patients. Data extracted from TCGA and visualised using cBioPortal (PanCancerAtlas). **B.** Dot plot map depicting scaled RNA expression of *FAM30A*, MSI2 and the stem cell markers CD34 and HOPX. Data extracted from^38^ and visualized using scRNAseq portal (Broad Institute). **C.** MSI2-CLIP experiment performed in NB4 cells^44^ and visualised using IGV software. Image is zoomed into the genomic localisation of the *FAM30A* locus.

**Supplementary Figure 5. Modulation of *FAM30A*/MSI2 differentially affects RUNX1 activation. A.** Schematic of regions where MSI2 binding was detected in AML and CML cells (CLIP experiments^42, 44^) in the RUNX1 isoforms. **B.** Quantitative analysis from Western blot of RUNX1A/B isoforms in stable KG-1a cell lines after 5 days in liquid culture. Normalisation was performed to control cells. **C.** Quantitative RT-qPCR analysis of RNA levels of RUNX1 isoforms in KG-1a cells under conditions as in B. **D**. Representative Western blot of stable MV4-11 cell lines. Normalisation was performed against control cells. Statistical significance was calculated using unpaired two-tailed student t-test. Data is displayed as mean values and error bars represent SEM.

**Supplementary Figure 6. Modulation of *FAM30A* expression alters leukemic engraftment in murine spleen. A.** Spleen weight (in mg) six weeks post-transplant of the stable KG-1a cell lines. Statistical significance was calculated using unpaired two-tailed student t-test. Data is displayed as mean values and error bars represent SEM.

## Supplementary Tables

**S1.** Cell lines used in this study.

**S2.** Plasmids and oligonucleotides used for cloning.

**S3.** Chemicals and reagents used in this study.

**S4.** RT-qPCR primers used in this study.

**S5.** Oligonucleotides used for *in vitro* transcription of biotinylated RNAs.

**S6.** Antibodies used in this study.

**S7.** RNA-sequencing results performed in stable KG-1a cell lines.

**S8.** GSEA analyses (Hallmarks) of the RNA-sequencing results in stable KG-1a cell lines.

**S9.** Proteomic analysis of significantly enriched proteins in pulldowns of the FAM30A repeats.

## Notes

### Competing Interest Statement

The authors have declared no competing interest.

## REFERENCES

1. Vetrie D, Helgason GV, Copland M. The leukaemia stem cell: similarities, differences and clinical prospects in CML and AML. Nat Rev Cancer 2020 Mar; 20(3): 158–173.

2. Misaghian N, Ligresti G, Steelman LS, Bertrand FE, Basecke J, Libra M, et al. Targeting the leukemic stem cell: the Holy Grail of leukemia therapy. Leukemia 2009 Jan; 23(1): 25–42.

3. Ng SWK, Murphy T, King I, Zhang T, Mah M, Lu Z, et al. A clinical laboratory-developed LSC17 stemness score assay for rapid risk assessment of patients with acute myeloid leukemia. Blood Adv 2022 Feb 8; 6(3): 1064–1073.

4. Vasseur L, Fenwarth L, Lambert J, de Botton S, Figeac M, Villenet C, et al. LSC17 score complements genetics and measurable residual disease in acute myeloid leukemia: an ALFA study. Blood Adv 2023 Aug 8; 7(15): 4024–4034.

5. Mercer TR, Dinger ME, Mattick JS. Long non-coding RNAs: insights into functions. Nat Rev Genet 2009 Mar; 10(3): 155–159.

6. Papaioannou D, Petri A, Dovey OM, Terreri S, Wang E, Collins FA, et al. The long non-coding RNA HOXB-AS3 regulates ribosomal RNA transcription in NPM1-mutated acute myeloid leukemia. Nat Commun 2019 Nov 25; 10(1): 5351.

7. Zhu G, Luo H, Feng Y, Guryanova OA, Xu J, Chen S, et al. HOXBLINC long non-coding RNA activation promotes leukemogenesis in NPM1-mutant acute myeloid leukemia. Nat Commun 2021 Mar 29; 12(1): 1956.

8. Liu S, Zhou J, Ye X, Chen D, Chen W, Lin Y, et al. A novel lncRNA SNHG29 regulates EP300-related histone acetylation modification and inhibits FLT3-ITD AML development. Leukemia 2023 Jul; 37(7): 1421–1434.

9. Tsai CH, Yao CY, Tien FM, Tang JL, Kuo YY, Chiu YC, et al. Incorporation of long non-coding RNA expression profile in the 2017 ELN risk classification can improve prognostic prediction of acute myeloid leukemia patients. EBioMedicine 2019 Feb; 40: 240–250.

10. Garzon R, Volinia S, Papaioannou D, Nicolet D, Kohlschmidt J, Yan PS, et al. Expression and prognostic impact of lncRNAs in acute myeloid leukemia. Proc Natl Acad Sci U S A 2014 Dec 30; 111(52): 18679–18684.

11. Bill M, Papaioannou D, Karunasiri M, Kohlschmidt J, Pepe F, Walker CJ, et al. Expression and functional relevance of long non-coding RNAs in acute myeloid leukemia stem cells. Leukemia 2019 Sep; 33(9): 2169–2182.

12. Luo H, Zhu G, Xu J, Lai Q, Yan B, Guo Y, et al. HOTTIP lncRNA Promotes Hematopoietic Stem Cell Self-Renewal Leading to AML-like Disease in Mice. Cancer Cell 2019 Dec 9; 36(6): 645–659 e648.

13. Al-Kershi S, Bhayadia R, Ng M, Verboon L, Emmrich S, Gack L, et al. The stem cell-specific long noncoding RNA HOXA10-AS in the pathogenesis of KMT2A-rearranged leukemia. Blood Adv 2019 Dec 23; 3(24): 4252–4263.

14. Metzeler KH, Hummel M, Bloomfield CD, Spiekermann K, Braess J, Sauerland MC, et al. An 86-probe-set gene-expression signature predicts survival in cytogenetically normal acute myeloid leukemia. Blood 2008 Nov 15; 112(10): 4193–4201.

15. Chuang MK, Chiu YC, Chou WC, Hou HA, Tseng MH, Kuo YY, et al. An mRNA expression signature for prognostication in de novo acute myeloid leukemia patients with normal karyotype. Oncotarget 2015 Nov 17; 6(36): 39098–39110.

16. Eppert K, Takenaka K, Lechman ER, Waldron L, Nilsson B, van Galen P, et al. Stem cell gene expression programs influence clinical outcome in human leukemia. Nat Med 2011 Aug 28; 17(9): 1086–1093.

17. Ng SW, Mitchell A, Kennedy JA, Chen WC, McLeod J, Ibrahimova N, et al. A 17-gene stemness score for rapid determination of risk in acute leukaemia. Nature 2016 Dec 15; 540(7633): 433–437.

18. Jung N, Dai B, Gentles AJ, Majeti R, Feinberg AP. An LSC epigenetic signature is largely mutation independent and implicates the HOXA cluster in AML pathogenesis. Nat Commun 2015 Oct 7; 6: 8489.

19. Elsayed AH, Rafiee R, Cao X, Raimondi S, Downing JR, Ribeiro R, et al. A six-gene leukemic stem cell score identifies high risk pediatric acute myeloid leukemia. Leukemia 2020 Mar; 34(3): 735–745.

20. Huang BJ, Smith JL, Farrar JE, Wang YC, Umeda M, Ries RE, et al. Integrated stem cell signature and cytomolecular risk determination in pediatric acute myeloid leukemia. Nat Commun 2022 Sep 19; 13(1): 5487.

21. Yang Y, Zhao Y, Hu N, Zhao J, Bai Y. lncRNA KIAA0125 functions as a tumor suppressor modulating growth and metastasis of colorectal cancer via Wnt/beta-catenin pathway. Cell Biol Int 2019 Dec; 43(12): 1463–1470.

22. Akrami S, Tahmasebi A, Moghadam A, Ramezani A, Niazi A. Integration of mRNA and protein expression data for the identification of potential biomarkers associated with pancreatic ductal adenocarcinoma. Comput Biol Med 2023 May; 157: 106529.

23. Wu G, Wang Q, Zhu T, Fu L, Li Z, Wu Y, et al. Identification and Validation of Immune-Related LncRNA Prognostic Signature for Lung Adenocarcinoma. Front Genet 2021; 12: 681277.

24. Hung SY, Lin CC, Hsu CL, Yao CY, Wang YH, Tsai CH, et al. The expression levels of long non-coding RNA KIAA0125 are associated with distinct clinical and biological features in myelodysplastic syndromes. Br J Haematol 2021 Feb; 192(3): 589–598.

25. Wang YH, Lin CC, Hsu CL, Hung SY, Yao CY, Lee SH, et al. Distinct clinical and biological characteristics of acute myeloid leukemia with higher expression of long noncoding RNA KIAA0125. Ann Hematol 2021 Feb; 100(2): 487–498.

26. Zhang T, Liao D, Hu Y. Cuproptosis-related lncRNAs forecast the prognosis of acute myeloid leukemia. Transl Cancer Res 2023 May 31; 12(5): 1175–1195.

27. de Lima DS, Cardozo LE, Maracaja-Coutinho V, Suhrbier A, Mane K, Jeffries D, et al. Long noncoding RNAs are involved in multiple immunological pathways in response to vaccination. Proc Natl Acad Sci U S A 2019 Aug 20; 116(34): 17121–17126.

28. Naldini MM, Casirati G, Barcella M, Rancoita PMV, Cosentino A, Caserta C, et al. Longitudinal single-cell profiling of chemotherapy response in acute myeloid leukemia. Nat Commun 2023 Mar 8; 14(1): 1285.

29. Haferlach T, Kohlmann A, Wieczorek L, Basso G, Kronnie GT, Bene MC, et al. Clinical utility of microarray-based gene expression profiling in the diagnosis and subclassification of leukemia: report from the International Microarray Innovations in Leukemia Study Group. J Clin Oncol 2010 May 20; 28(15): 2529–2537.

30. Tang Z, Kang B, Li C, Chen T, Zhang Z. GEPIA2: an enhanced web server for large-scale expression profiling and interactive analysis. Nucleic Acids Res 2019 Jul 2; 47(W1): W556–W560.

31. Valk PJ, Verhaak RG, Beijen MA, Erpelinck CA, Barjesteh van Waalwijk van Doorn-Khosrovani S, Boer JM, et al. Prognostically useful gene-expression profiles in acute myeloid leukemia. N Engl J Med 2004 Apr 15; 350(16): 1617–1628.

32. Herold T, Jurinovic V, Batcha AMN, Bamopoulos SA, Rothenberg-Thurley M, Ksienzyk B, et al. A 29-gene and cytogenetic score for the prediction of resistance to induction treatment in acute myeloid leukemia. Haematologica 2018 Mar; 103(3): 456–465.

33. Verhaak RG, Wouters BJ, Erpelinck CA, Abbas S, Beverloo HB, Lugthart S, et al. Prediction of molecular subtypes in acute myeloid leukemia based on gene expression profiling. Haematologica 2009 Jan; 94(1): 131–134.

34. Raponi M, Lancet JE, Fan H, Dossey L, Lee G, Gojo I, et al. A 2-gene classifier for predicting response to the farnesyltransferase inhibitor tipifarnib in acute myeloid leukemia. Blood 2008 Mar 1; 111(5): 2589–2596.

35. Papaemmanuil E, Gerstung M, Bullinger L, Gaidzik VI, Paschka P, Roberts ND, et al. Genomic Classification and Prognosis in Acute Myeloid Leukemia. N Engl J Med 2016 Jun 9; 374(23): 2209–2221.

36. Li K, Du Y, Cai Y, Liu W, Lv Y, Huang B, et al. Single-cell analysis reveals the chemotherapy-induced cellular reprogramming and novel therapeutic targets in relapsed/refractory acute myeloid leukemia. Leukemia 2023 Feb; 37(2): 308–325.

37. Boutzen H, Madani Tonekaboni SA, Chan-Seng-Yue M, Murison A, Takayama N, Mbong N, et al. A primary hierarchically organized patient-derived model enables in depth interrogation of stemness driven by the coding and non-coding genome. Leukemia 2022 Nov; 36(11): 2690–2704.

38. Lasry A, Nadorp B, Fornerod M, Nicolet D, Wu H, Walker CJ, et al. An inflammatory state remodels the immune microenvironment and improves risk stratification in acute myeloid leukemia. Nat Cancer 2023 Jan; 4(1): 27–42.

39. Zhang K, Shi ZM, Chang YN, Hu ZM, Qi HX, Hong W. The ways of action of long non-coding RNAs in cytoplasm and nucleus. Gene 2014 Aug 15; 547(1): 1–9.

40. Duggimpudi S, Kloetgen A, Maney SK, Munch PC, Hezaveh K, Shaykhalishahi H, et al. Transcriptome-wide analysis uncovers the targets of the RNA-binding protein MSI2 and effects of MSI2’s RNA-binding activity on IL-6 signaling. J Biol Chem 2018 Oct 5; 293(40): 15359–15369.

41. Nguyen DTT, Lu Y, Chu KL, Yang X, Park SM, Choo ZN, et al. HyperTRIBE uncovers increased MUSASHI-2 RNA binding activity and differential regulation in leukemic stem cells. Nat Commun 2020 Apr 24; 11(1): 2026.

42. Park SM, Gonen M, Vu L, Minuesa G, Tivnan P, Barlowe TS, et al. Musashi2 sustains the mixed-lineage leukemia-driven stem cell regulatory program. J Clin Invest 2015 Mar 2; 125(3): 1286–1298.

43. Wesely J, Kotini AG, Izzo F, Luo H, Yuan H, Sun J, et al. Acute Myeloid Leukemia iPSCs Reveal a Role for RUNX1 in the Maintenance of Human Leukemia Stem Cells. Cell Rep 2020 Jun 2; 31(9): 107688.

44. Rentas S, Holzapfel N, Belew MS, Pratt G, Voisin V, Wilhelm BT, et al. Musashi-2 attenuates AHR signalling to expand human haematopoietic stem cells. Nature 2016 Apr 28; 532(7600): 508–511.

45. Schwarzer A, Emmrich S, Schmidt F, Beck D, Ng M, Reimer C, et al. The non-coding RNA landscape of human hematopoiesis and leukemia. Nat Commun 2017 Aug 9; 8(1): 218.

46. Di Giacomo D, La Starza R, Gorello P, Pellanera F, Kalender Atak Z, De Keersmaecker K, et al. 14q32 rearrangements deregulating BCL11B mark a distinct subgroup of T-lymphoid and myeloid immature acute leukemia. Blood 2021 Sep 2; 138(9): 773–784.

47. Onoguchi M, Zeng C, Matsumaru A, Hamada M. Binding patterns of RNA-binding proteins to repeat-derived RNA sequences reveal putative functional RNA elements. NAR Genom Bioinform 2021 Sep; 3(3): lqab055.

48. McHugh CA, Chen CK, Chow A, Surka CF, Tran C, McDonel P, et al. The Xist lncRNA interacts directly with SHARP to silence transcription through HDAC3. Nature 2015 May 14; 521(7551): 232–236.

49. Nagase T, Seki N, Tanaka A, Ishikawa K, Nomura N. Prediction of the coding sequences of unidentified human genes. IV. The coding sequences of 40 new genes (KIAA0121-KIAA0160) deduced by analysis of cDNA clones from human cell line KG-1. DNA Res 1995 Aug 31; 2(4): 167–174, 199-210.

50. Sun L, Wang W, Han C, Huang W, Sun Y, Fang K, et al. The oncomicropeptide APPLE promotes hematopoietic malignancy by enhancing translation initiation. Mol Cell 2021 Nov 4; 81(21): 4493–4508 e4499.

51. de Andres-Aguayo L, Varas F, Kallin EM, Infante JF, Wurst W, Floss T, et al. Musashi 2 is a regulator of the HSC compartment identified by a retroviral insertion screen and knockout mice. Blood 2011 Jul 21; 118(3): 554–564.

52. Park SM, Deering RP, Lu Y, Tivnan P, Lianoglou S, Al-Shahrour F, et al. Musashi-2 controls cell fate, lineage bias, and TGF-beta signaling in HSCs. J Exp Med 2014 Jan 13; 211(1): 71–87.

53. Zhu Y, Zhou B, Hu X, Ying S, Zhou Q, Xu W, et al. LncRNA LINC00942 promotes chemoresistance in gastric cancer by suppressing MSI2 degradation to enhance c-Myc mRNA stability. Clin Transl Med 2022 Jan; 12(1): e703.

54. Zhu P, He F, Hou Y, Tu G, Li Q, Jin T, et al. A novel hypoxic long noncoding RNA KB-1980E6.3 maintains breast cancer stem cell stemness via interacting with IGF2BP1 to facilitate c-Myc mRNA stability. Oncogene 2021 Mar; 40(9): 1609–1627.

55. Wang Y, Zhu P, Luo J, Wang J, Liu Z, Wu W, et al. LncRNA HAND2-AS1 promotes liver cancer stem cell self-renewal via BMP signaling. EMBO J 2019 Sep 2; 38(17): e101110.

56. Kharas MG, Lengner CJ, Al-Shahrour F, Bullinger L, Ball B, Zaidi S, et al. Musashi-2 regulates normal hematopoiesis and promotes aggressive myeloid leukemia. Nat Med 2010 Aug; 16(8): 903–908.

57. Vu LP, Prieto C, Amin EM, Chhangawala S, Krivtsov A, Calvo-Vidal MN, et al. Functional screen of MSI2 interactors identifies an essential role for SYNCRIP in myeloid leukemia stem cells. Nat Genet 2017 Jun; 49(6): 866–875.

58. Minuesa G, Albanese SK, Xie W, Kazansky Y, Worroll D, Chow A, et al. Small-molecule targeting of MUSASHI RNA-binding activity in acute myeloid leukemia. Nat Commun 2019 Jun 19; 10(1): 2691.

59. North TE, de Bruijn MF, Stacy T, Talebian L, Lind E, Robin C, et al. Runx1 expression marks long-term repopulating hematopoietic stem cells in the midgestation mouse embryo. Immunity 2002 May; 16(5): 661–672.

60. Liu X, Zhang Q, Zhang DE, Zhou C, Xing H, Tian Z, et al. Overexpression of an isoform of AML1 in acute leukemia and its potential role in leukemogenesis. Leukemia 2009 Apr; 23(4): 739–745.

61. Gialesaki S, Brauer-Hartmann D, Issa H, Bhayadia R, Alejo-Valle O, Verboon L, et al. RUNX1 isoform disequilibrium promotes the development of trisomy 21-associated myeloid leukemia. Blood 2023 Mar 9; 141(10): 1105–1118.

62. Ghozi MC, Bernstein Y, Negreanu V, Levanon D, Groner Y. Expression of the human acute myeloid leukemia gene AML1 is regulated by two promoter regions. Proc Natl Acad Sci U S A 1996 Mar 5; 93(5): 1935–1940.

63. Navarro-Montero O, Ayllon V, Lamolda M, Lopez-Onieva L, Montes R, Bueno C, et al. RUNX1c Regulates Hematopoietic Differentiation of Human Pluripotent Stem Cells Possibly in Cooperation with Proinflammatory Signaling. Stem Cells 2017 Nov; 35(11): 2253–2266.

64. Hornung R, Jurinovic V, Batcha AMN, Bamopoulos SA, Rothenberg-Thurley M, Amler S, et al. Mediation analysis reveals common mechanisms of RUNX1 point mutations and RUNX1/RUNX1T1 fusions influencing survival of patients with acute myeloid leukemia. Sci Rep 2018 Jul 26; 8(1): 11293.

65. Salem M, Delwel R, Touw I, Mahmoud LA, Elbasousy EM, Lowenberg B. Modulation of colony stimulating factor-(CSF) dependent growth of acute myeloid leukemia by tumor necrosis factor. Leukemia 1990 Jan; 4(1): 37–43.

66. Yoshimoto G, Miyamoto T, Jabbarzadeh-Tabrizi S, Iino T, Rocnik JL, Kikushige Y, et al. FLT3-ITD up-regulates MCL-1 to promote survival of stem cells in acute myeloid leukemia via FLT3-ITD-specific STAT5 activation. Blood 2009 Dec 3; 114(24): 5034–5043.

67. Wisniewski JR, Zougman A, Nagaraj N, Mann M. Universal sample preparation method for proteome analysis. Nat Methods 2009 May; 6(5): 359–362.

68. Kim D, Langmead B, Salzberg SL. HISAT: a fast spliced aligner with low memory requirements. Nat Methods 2015 Apr; 12(4): 357–360.

69. Liao Y, Smyth GK, Shi W. featureCounts: an efficient general purpose program for assigning sequence reads to genomic features. Bioinformatics 2014 Apr 1; 30(7): 923–930.

70. Yates AD, Achuthan P, Akanni W, Allen J, Alvarez-Jarreta J, Amode MR, et al. Ensembl 2020. Nucleic Acids Res 2020 Jan 8; 48(D1): D682–D688.

71. Robinson MD, McCarthy DJ, Smyth GK. edgeR: a Bioconductor package for differential expression analysis of digital gene expression data. Bioinformatics 2010 Jan 1; 26(1): 139–140.

72. Wallace IM, O’Sullivan O, Higgins DG, Notredame C. M-Coffee: combining multiple sequence alignment methods with T-Coffee. Nucleic Acids Res 2006; 34(6): 1692–1699.

73. Subramanian A, Tamayo P, Mootha VK, Mukherjee S, Ebert BL, Gillette MA, et al. Gene set enrichment analysis: a knowledge-based approach for interpreting genome-wide expression profiles. Proc Natl Acad Sci U S A 2005 Oct 25; 102(43): 15545–15550.

74. Rapin N, Bagger FO, Jendholm J, Mora-Jensen H, Krogh A, Kohlmann A, et al. Comparing cancer vs normal gene expression profiles identifies new disease entities and common transcriptional programs in AML patients. Blood 2014 Feb 6; 123(6): 894–904.

