## Supplementary Figures for "The long non-coding RNA *FAM30A* regulates the Musashi2-RUNX1 axis and is required for LSC function in AML cells"

### Slide 1
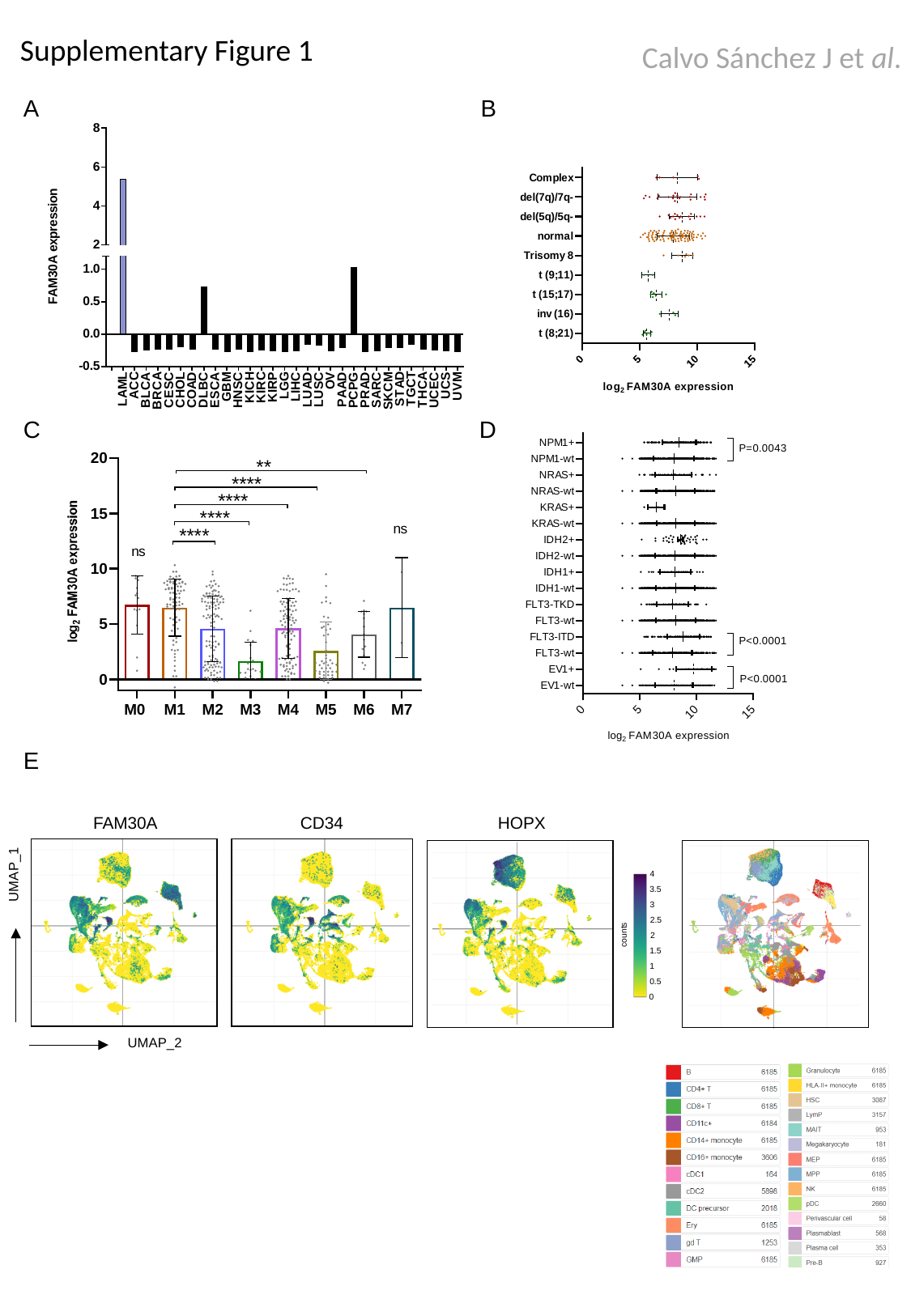

Supplementary Figure 1
Calvo Sánchez J et al.
A
B
C
D
E
FAM30A
CD34
HOPX
UMAP_1
UMAP_2
4
3.5
3
2.5
counts
2
1.5
1
0.5
0

### Slide 2
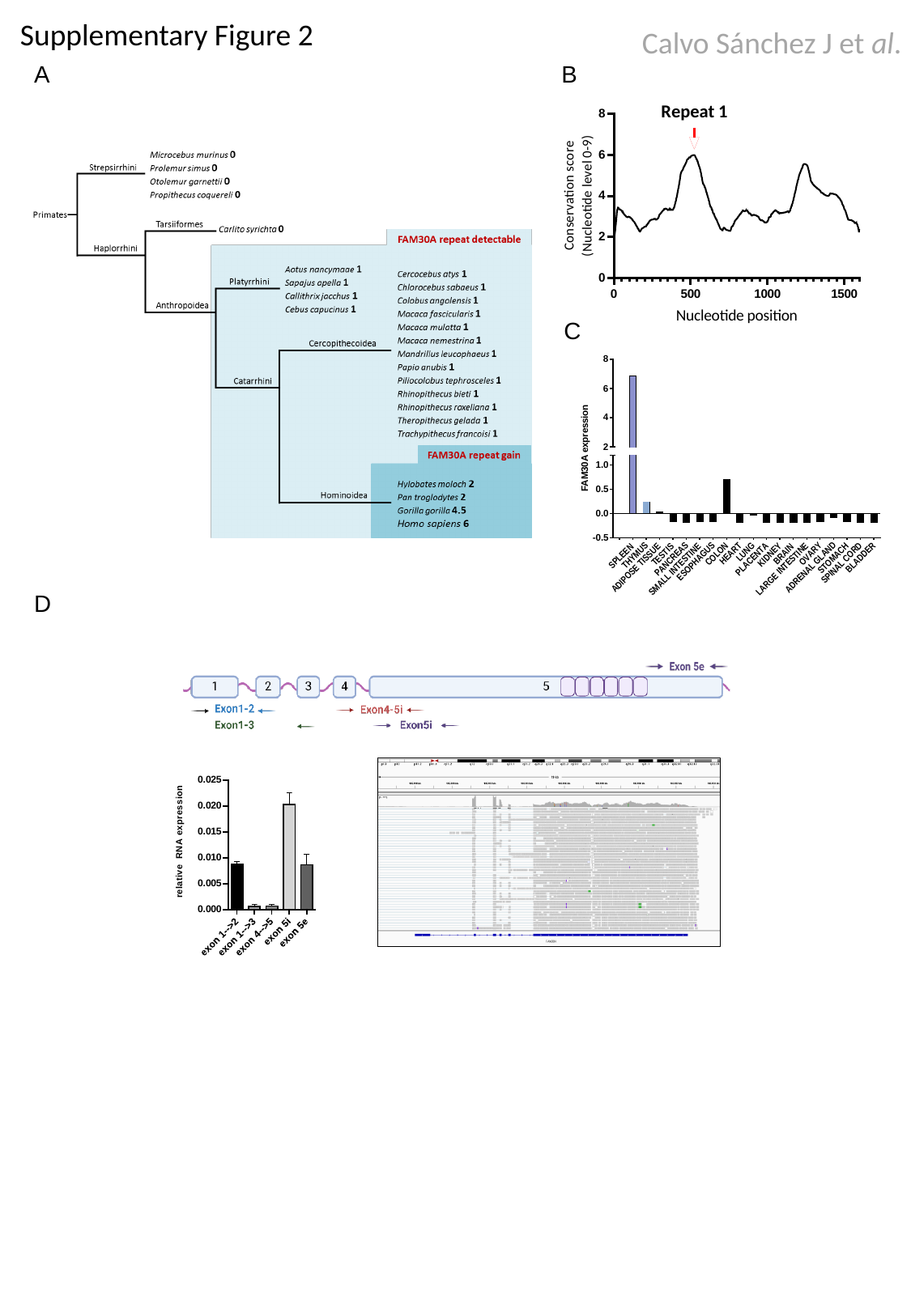

Supplementary Figure 2
Calvo Sánchez J et al.
A
B
C
D

### Slide 3
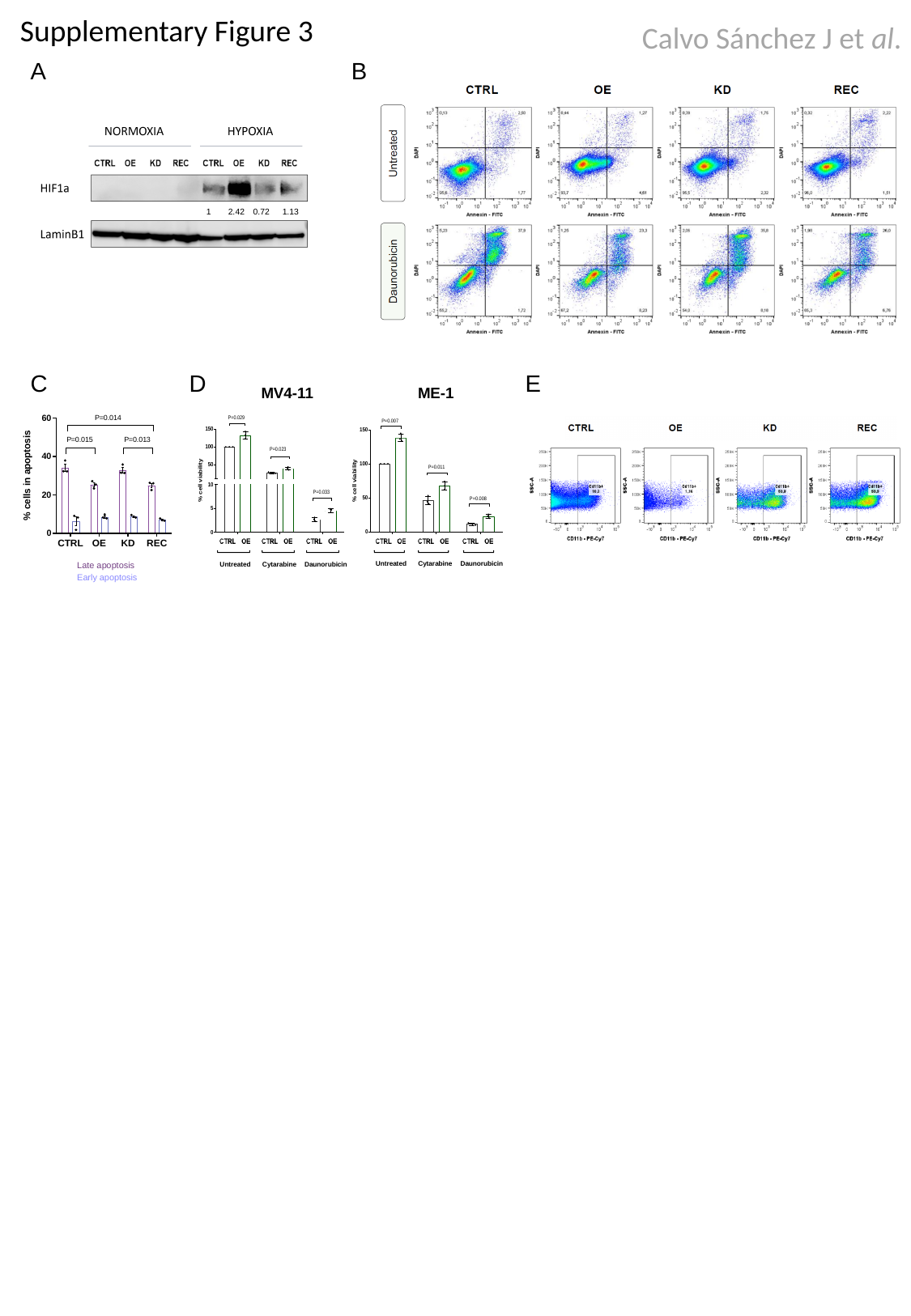

Supplementary Figure 3
Calvo Sánchez J et al.
A
B
1
2.42
0.72
1.13
C
D
E
MV4-11
ME-1
Untreated
Cytarabine
Daunorubicin
Cytarabine
Daunorubicin
Untreated

### Slide 4
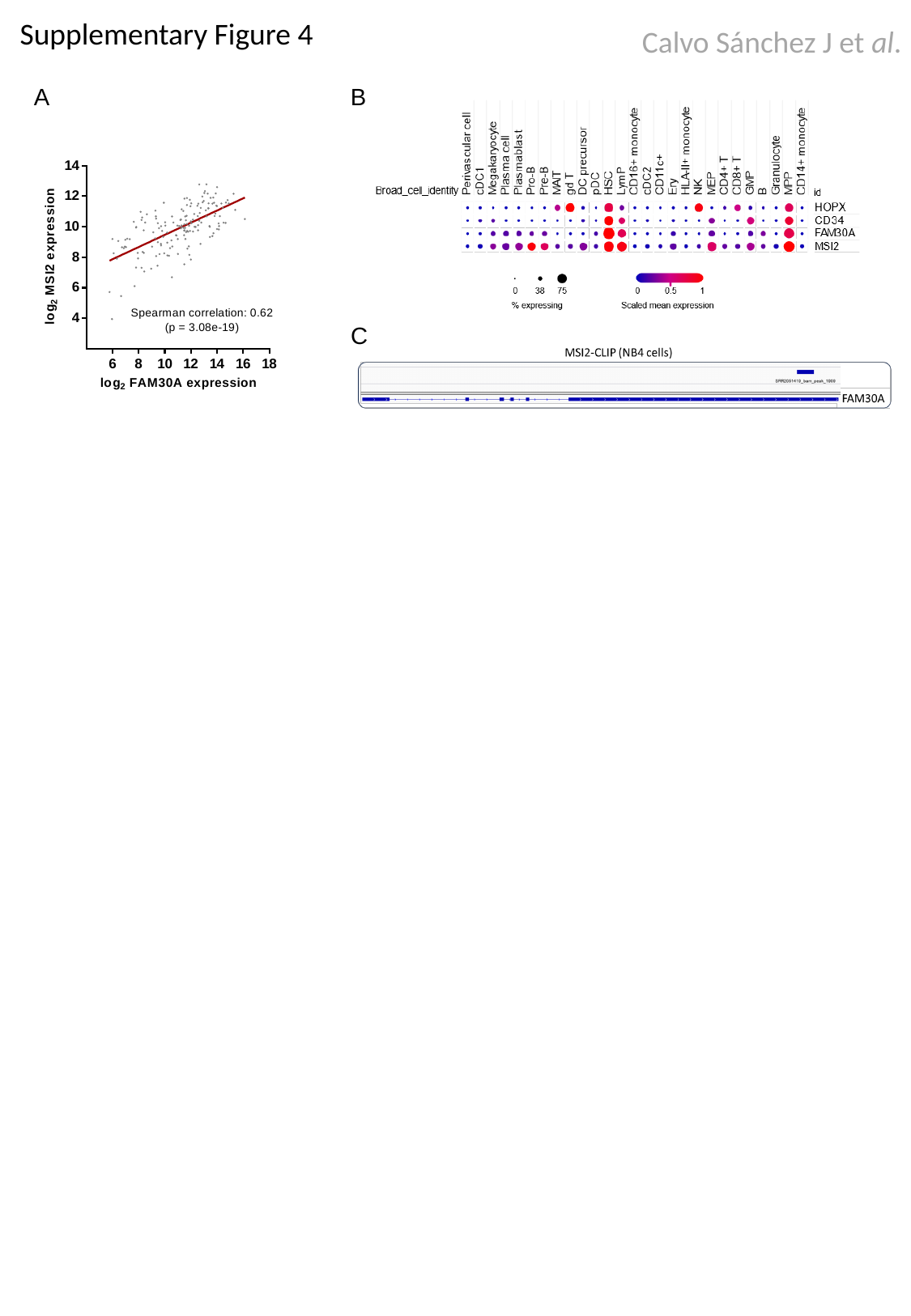

Supplementary Figure 4
Calvo Sánchez J et al.
A
B
C

### Slide 5
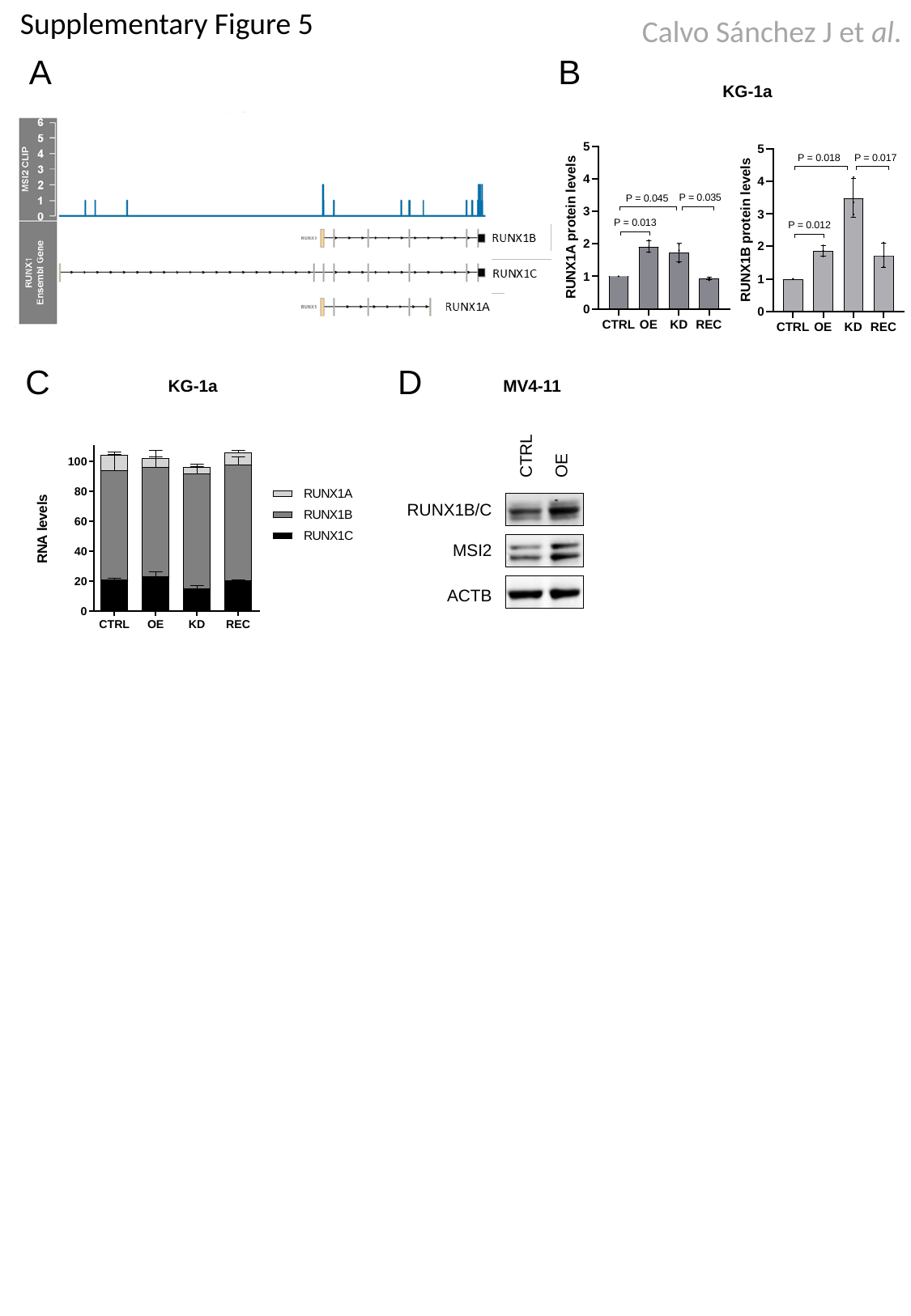

Supplementary Figure 5
Calvo Sánchez J et al.
A
B
KG-1a
C
D
CTRL
OE
RUNX1B/C
MSI2
ACTB
KG-1a
MV4-11

### Slide 6
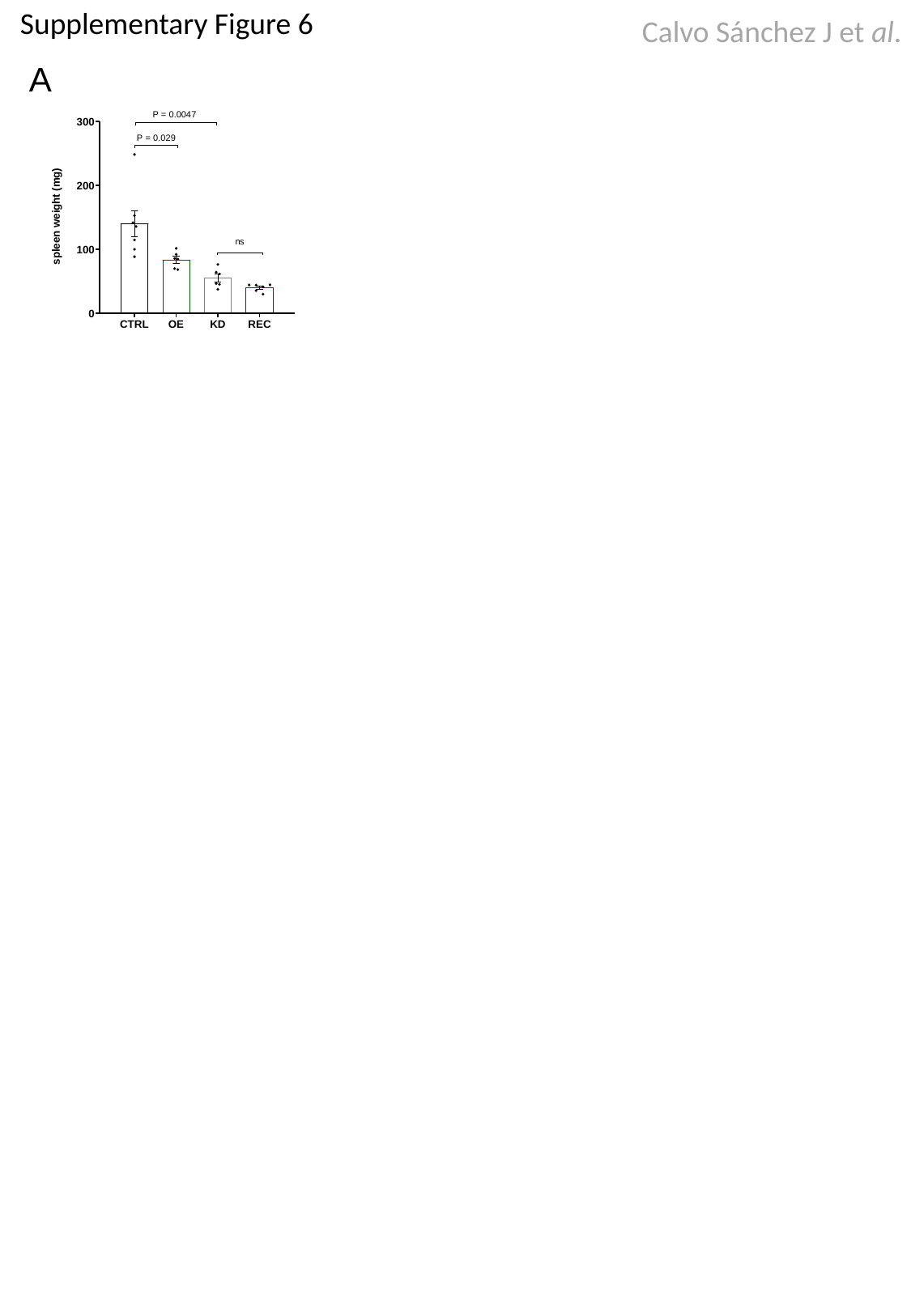

Supplementary Figure 6
Calvo Sánchez J et al.
A
